# Biophysical Basis of the *in vivo* Electroretinogram of the Mouse: Current Source Density Analysis of Genetically and Pharmacologically Isolated Rod photoreceptor-driven Currents

**DOI:** 10.1101/2025.02.05.636690

**Authors:** Gabriel Peinado Allina, Prescott Alexander, Kaitryn Ronning, Marie E. Burns, Edward N. Pugh

## Abstract

To better understand the molecular basis of the mouse electroretinogram (ERG) we have developed a biophysical model of the rod photoreceptor layer’s ionic mechanisms and applied current source-density (CSD) analysis to predict the genetically and pharmacologically isolated rod ERG *a*-wave. The saturating *a*-wave is characterized by a rapid relaxation (τ ~ 6 ms) from its maximum that has been hypothesized to be caused by HCN1 channel opening consequent to light-triggered sustained hyperpolarization, or by extracellular flow of capacitive current during hyperpolarization. To test these hypotheses, the CSD model included an ensemble of 11 rods with cell body locations spanning the outer nuclear layer, and with ionic mechanisms -- including CNG channels, NCKX in the outer segment, NKX, Kv2.1 channels in the inner segment, and HCN1 throughout the “non-outer segment” -- fully specified as to axial distributions, voltage dependencies, and the extracellular conductivity of the photoreceptor layer extracellular space. Predicting with CSD the steady-state axial distributions of rod dark current and transretinal potential, and the spatio-temporal response of the activation phase of the rod’s light response to intense stimuli, the analysis confirmed the Robson-Frishman hypothesis that extracellular capacitive current flowing towards the inner segment upon the rapid closure of CNG current plays a major role in shaping the early *a*-wave. The CSD analysis also reveals that extracellular current from HCN1 channels opened by the hyperpolarization contributes a net extracellular current flowing toward the ONL that contributes materially to the saturated *a*-wave relaxation and to setting its plateau level.

## INTRODUCTION

This, the translational era of biomedical science, has an imperative to integrate advances in characterization of molecular structure and function obtained *ex vivo* with cellular and tissue function *in vivo*. This imperative arises both from the ultimate scientific goal of integrating biomedical science from the molecular level to that of the whole organism, and from the practical goal providing medicine with interpretable measures for assessing health and disease status and the efficaciousness of therapy. To achieve these goals, biomedical science and physiology in particular need to evolve “back to the future”, reintegrating classic biophysically rationalized tissue- and organ-level analyses with more recent cellular and molecular mechanistic insight.

The retina is a portion of the central nervous system that is remarkably accessible for *in vivo* electrophysiological (Perlman, 1995; Thomas and Lamb, 1999; Robson et al., 2004; Kenkre et al., 2005; Robson and Frishman, 2014) and optophysiological (Hillmann et al., 2016; Zhang et al., 2017; Azimipour et al., 2020; Cooper et al., 2020; Pandiyan et al., 2020; Pandiyan et al., 2022) measurements in both humans and experimental animals. In the 55 years since the seminal discovery of the dark current of rods and its suppression by light (Hagins et al., 1970a; Penn and Hagins, 1972), a vast body of knowledge has accrued about the biochemistry, physiology and molecular biology of retinal photoreceptors, including their ion channel expression and the molecular mechanisms of phototransduction, and about the structure and function of supporting ocular cells and tissues. The value of this accrued knowledge for translational medicine of the eye would be considerably enhanced were it fully incorporated into a biophysical analysis of the field potentials generated by photoreceptor responses.

The physiological function of electrically active cells *in vivo* is affected by their morphology, their ion channel expression patterns and the properties of the tissues in which they reside. As the proportionality factor between the local extracellular current density and its field potential, accurately determined conductivity is critical for current source density (CSD) analysis, which seeks to quantitatively link measured field potentials to their underlying cellular current generators. For field potentials originating in the photoreceptor layer of the retina, these generators include not only the radially distributed currents of rod and cone ion channels and transporters, but also radially flowing capacitive currents. (Robson and Frishman, 2014), and transepithelial currents arising from light-dependent changes of K^+^ _o_ in the subretinal space (Steinberg et al., 1980; Steinberg, 1985).

In this investigation we take advantage of recent advances in characterization of the molecular mechanisms underlying the dark current of mouse rods and their expression patterns (Fortenbach *et al*, 2021), in the development of genetic (Ronning et al., 2018) and pharmacological tools for isolating rod photoreceptor components of the corneal electroretinogram (ERG) (reviewed in (Robson and Frishman, 2014), and undertake to more completely quantify than heretofore the relationship between rod photoreceptor ionic mechanisms and the field potentials to which they give rise *in vivo*.

## THEORY

### Background, rationale and objective

Hagins *et al*. (1970) discovered the rod photoreceptor dark current by performing a classic current source density (CSD) analysis of the photoreceptor layer of rat retinal slices. Their analysis applied the Equation of Continuity (EOC),

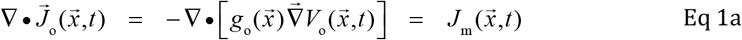

to extracellular potentiometric data obtained at radial positions throughout the photoreceptor layer to deduce the distribution of current sources and sinks. In Eq 1a 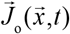 is the current density vector (A cm^−2^), *V*_*o*_(*x, y, z*) the extracellular electrical potential at each location in the tissue, *g*_o_ (S cm^−1^), a 3×3 position-dependent conductivity tensor, ▽ • and 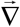 the divergence and gradient operators respectively, and *J*_m_ (*x, y, z*) (A cm^−3^) the source/sink volume current density. The leftmost term of Eq 1a is the divergence of the three orthogonal components of the local current density in the volume element, while the middle term embodies the insight that the current density vector can be expressed as the matrix product of the negative gradient of a potential and the conductivity when magnetic fields are negligible. The rightmost term of Eq 1a must be zero unless charge is flowing into or out of the extracellular volume element, and thus deviations from zero are assigned to local cellular (transmembrane) sources or sinks, respectively. Because of the transverse symmetry of the retinal slice, Hagins *et al*. (1970) reduced Eq 1a to the corresponding expression with only one spatial coordinate, i.e.,

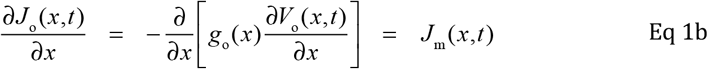

where *x* is radial (depth) position in the retina, *J*_*o*_(*x, t*) are *V*_*o*_(*x, t*) are the extracellular current density and potential respectively, *g*_o_ (*x*) (S cm^−1^) the extracellular conductivity density and *J*_m_ (A cm^−3^) the volume density of all membrane (ionic and capacitive) current contributing as source or sink to the specific volume element.

Hagins *et al*. (1970) used Eq 1b in the “inverse” direction, that is, applied it to measurements of *V*_o_ and *g*_o_ to deduce *J*_m_ (*x,t*), the axial distribution of membrane current sources and sinks. A primary objective of the present paper is to apply CSD analysis (Eq 1b) in the “forward” direction. Specifically, the goal has been to develop a biophysically accurate description of the axial distribution of membrane currents of rod photoreceptors and the conductivity profile of the photoreceptor layer in the mouse retina, to use this information to predict the extracellular potential *V*_*o*_(*x, t*) in darkness and in response to light, and finally to compare these predictions with the genetically and pharmacologically isolated photoreceptor current components of the corneal ERG measured *in vivo*.

Developing a biophysically accurate “forward application” of Eq 1b to predict field potentials arising from the photoreceptor layer of the retina is challenging for several reasons, including uncertainty about the molecular identity and precise axial distribution of relevant ion channels and electrogenic transporters, and the theoretical and computational problems arising from cross-coupling between photoreceptors via their contributions to *V*_o_. While recent work characterizing the ionic mechanisms of mouse rods (Fortenbach *et al*., 2021) and data presented here allow us to address most of these issues, theoretical treatment of the cross-coupling between photoreceptors is notably challenging. To appreciate why, consider an analysis of the total volume current density *J*_m_ (*x,t*) (units: A cm^−3^) of membrane current (Eq 1b) into the ionic and capacitive contributions at depth *x* of an appropriate ensemble of *N*_R_ morphologically distinct rod types (Figs. 1, 2):

**Figure 1.**
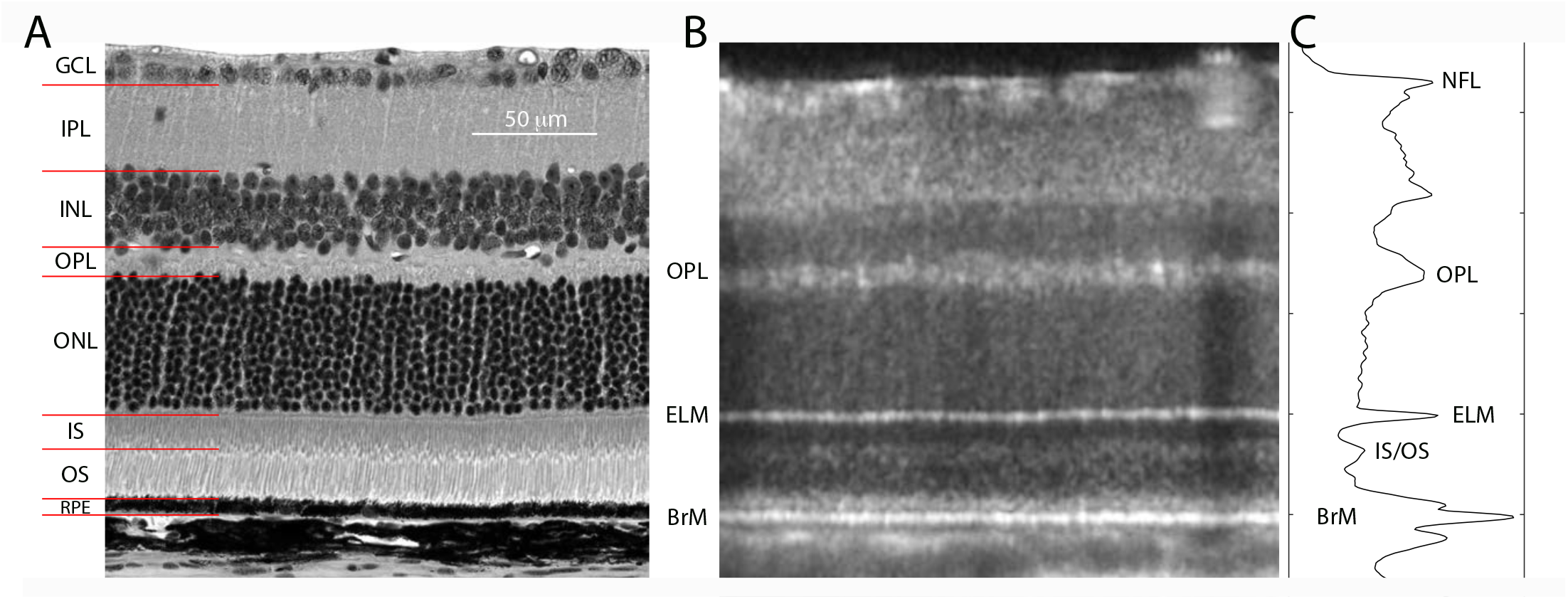
Spatial dimensions of the mouse photoreceptor layer. **A**. Transmitted light microscopic image of a thin section of C57Bl6 mouse. **B**. Optical coherence tomographic “B-scan” of a live (albino) mouse retina. **C**. Profile plot of the spatially averaged scattering intensity with the principal scattering bands identified. Abbreviations: NFL, neurofibrillary layer; GCL, ganglion cell layer; IPL, inner plexiform layer; INL, inner nuclear layer; OPL, outer plexiform layer; ONL, outer nuclear layer; IS, inner segment layer; OS, outer segment layer; RPE, retinal pigment epithelium; ELM, external limiting membrane; IS/OS, inner segment – outer segment transition zone; BrM, Bruch’s membrane (RPE basement membrane).

**Figure 2.**
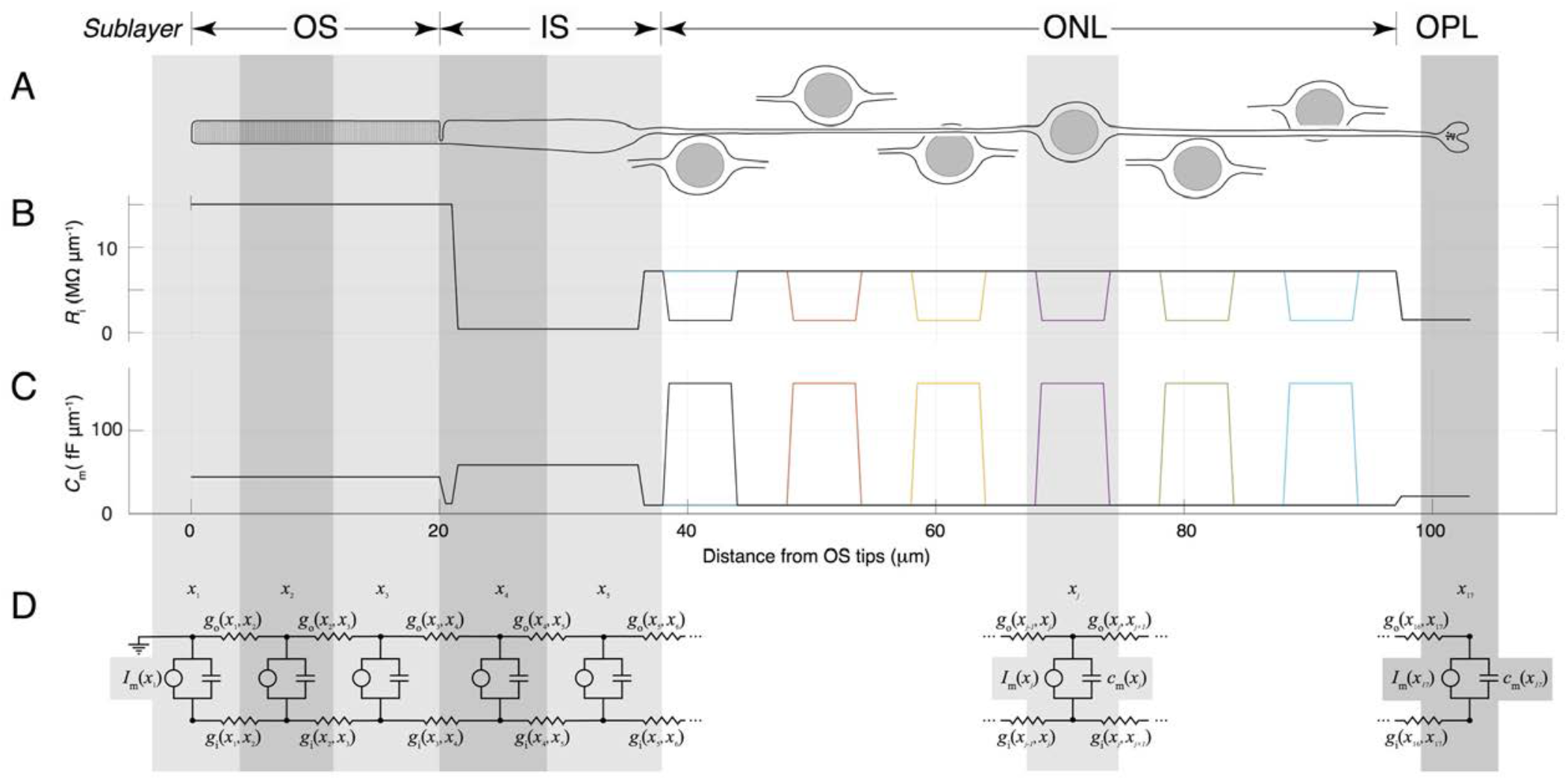
Spatial and static electrical features of the rod ensemble model of the photoreceptor layer. **A**. Schematic drawing of a single mouse rod and cell bodies of other rods. **B**. The internal axial resistance profile of 6 of the 11 mouse rods comprising the ensemble (all 11 model rods are assumed to have the same OS and IS dimensions and positioning; only 6 are shown for clarity). **C**. Capacitance profile of the same set of 6 rods. D. Classic cable electrical schematic of a single model rod discretized into segments bracketed by the axial positions *x*_1_ = 0, …, *x*_Nx_. (See text for further details.)

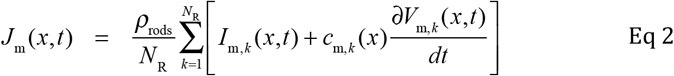

Here ρ_rods_ the spatial density of rods (rods cm^−2^) and *k* indexes the distinct cell types of the ensemble, with *I*_m, *k*_ (*x,t*)(A cm^−1^) the axial density of membrane current and *c*_m, *k*_ (*x*) (F cm^−1^) the axial capacitance density of the *k*^th^ rod type of the ensemble. The local membrane potential *V*_m, *k*_ (*x,t*) of the *k*^th^ cell type necessarily depends on the local external potential *V*_o_(*x, t*), which in turn depends on the ionic and capacitive currents of *all* the cells that contribute current to the extracellular space surrounding cells of type *k*. Specifically, both the steady state membrane potential distribution and the dynamic electrical activity of such an ensemble during the light response are cross-coupled through their mutual contributions to *V*_o_(*x, t*), the local instantaneous extracellular potential. (Because the rod spatial density ρ_rods_ does not differentiate amongst rod types, the “average rod” of the ensemble serves as the unit with respect to spatial density.)

Such cross-coupling between cells would be immaterial were the cells in the photoreceptor layer homogeneous, i.e., essentially identical. One violation is the presence of cone photoreceptors, which comprise 3% of mouse photoreceptors. A second violation of cellular homogeneity is the axially distributed locations of rod cell bodies across the outer nuclear layer (ONL) (Fig. 1A). The distribution of the cell bodies over the ONL predictably produces systematic variation amongst rods in their ionic and capacitive contributions to the transretinal photovoltage (Robson and Frishman, 2014). A third violation of homogeneity arises from the presence of Müller glia, which span the retina from the ELM to the NFL, and also have processes that reach well into the IS and even into the OS layer (Fig. 1). Müller glia not only are present in most of the photoreceptor layer, but also play an active role in management of K^+^_o_ in the subretinal space (Steinberg et al., 1980; Steinberg, 1985), a restricted extracellular volume between the ELM and the RPE (Fig. 1), which forms the posterior blood-retinal barrier. Even neglecting light- and time-varying contributions arising from cones and Müller glia, the structural variation amongst rods presents a problem for predicting *V*_*o*_(*x, t*). Here we address this latter variation, generalizing the single-rod models of Hagins *et al*. (1970) and of (Robson and Frishman, 2014) to an ensemble of rods with cell body positions spanning the ONL.

### Defining the rod ensemble

To predict the trans-photoreceptor layer potential in the dark and in response to light the following spatio-temporal differential equation for *V*_m, *k*_ (*x,t*) and *V*_o_ (*x,t*) must be solved, subject to appropriate boundary and initial conditions, and a model of the light-driven suppression of outer segment CNG current:

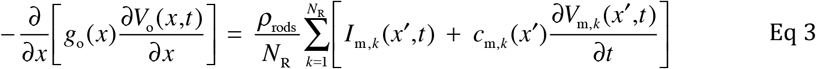

Equation 3 is derived by substitution into Eq 1b of the expression for *J*_m_(*x,t*) in Eq 2. To completely define the ensemble model embodied in Eq 3, a number of properties of the photoreceptor layer and the rods must be specified, including the external conductivity profile *g*_o_ (*x*), the molecular identities and other features of the ionic mechanisms underlying the membrane currents *I*_m, *k*_ (*x,t*), including their voltage-dependencies, transition kinetics and axial distributions in the different rod types, and the internal resistance and the capacitance distributions *c*_m, *k*_ (*x*) of each rod of the ensemble. Tables 1 and 2 provide a summary of relevant properties of mouse rods, while Figure 2 serves to illustrate graphically how the ensemble was defined. The immediately following paragraphs provide further detail and explanations.

**Table 1.** Structural Features of Mouse Rod Photoreceptor and the Photoreceptor Layer.

| Retinal Layer | Thickness/<br>length | Diameter | Axial Resistance<sup>a</sup> | Capacitance | Refs | Description/Comment |
| --- | --- | --- | --- | --- | --- | --- |
| Name/abbrev. | μm | μm | MΩ μm<sup>-1</sup> | fF μm<sup>-1</sup> |  |  |
| Outer segment | 20 | 1.4 | 15 | 44.0 | (1) | 5% patent cross section:<br>cytoplasmic gap (15 nm) of disc |
| Connecting cilium | 1 | 2.0 | 15 | 12 |  |  |
| Inner segment | 15 | 1.9 | 0.42 | 58.6 |  | IS layer packing fraction: 0.92 |
| Axon | 61 | 0.45 | 7.25 | 10 | (1) | Length includes cell body |
| Cell body | -- | 5.0 | 1.45 | 157 | Fig. 1A | <sup>b</sup>Size constrained by ONL packing;<br>resistance by shell of cytoplasm |
| Synaptic layer<br>(OPL) | 6.0 |  |  | 21 | (2) | <sup>c</sup>Size constrained by OPL packing<br>of rod spherules and cone pedicles |

*Table notes.* <sup>a</sup>Defined by $R_l(x) = \rho / A(x)$ where $\rho$ is cytoplasm resistivity (taken to be 115 Ω cm (Fry et al., 2012)), and $A(x)$ is the cross-sectional area open or patent to axial current flow. Total capacitance of each rod 3.3 pF.
*Table references.* <sup>1</sup>(Carter-Dawson and LaVail, 1979) <sup>2</sup>(Li et al., 2016)

**Table 2.** Variables, parameters and their units of the rod ensemble model.

| *Variable* | *Unit* | *Comment* | *Equations* | *Figures* |
| --- | --- | --- | --- | --- |
| $t$ | s | $t = 0$ corresponds to time of a brief (1 ms) light flash | 3 | 6–8 |
| $x$ | cm | retinal depth re. rod tips ($x=0$); converted to $\mu\text{m}$ in figures; | | 2–5; |
| $V_o(x,t)$ | V | Trans-photoreceptor layer (PRL) potential; “o” = “outside” | 3, 21b | 7B, C |
| $J_o(x,t)$ | A cm<sup>-2</sup> | extracellular current density in PRL at depth $x$ | 7, 12 | cf. S1 |
| $V_{i,k}(x,t)$ | V | Internal axial potential of rod $k$ of ensemble; “i” = “inside” | 10, 23b | – |
| $j_{i,k}(x,t)$ | A | current flowing internally in rod $k$ at PRL depth $x$ | 9, 10 | – |
| $I_{m,k}(x,t)$ | A cm<sup>-1</sup> | ionic membrane current density of rod $k$ at PRL depth $x$ | 2, 4 | cf. 7E, F |
| $V_{m,k}(x,t)$ | V | transmembrane potential of rod $k$ at PRL depth $x$ | 2, 11 | 5, 7D |
| *Parameter* | *Unit* |  |  |  |
| $L$ | cm | Length/depth of photoreceptor layer (PRL) | 4 | Figs. 1, 2 |
| $\rho_{\text{rods}}$ | cm<sup>-2</sup> | Rod spatial density | 2, 3, 12, 13b | |
| $g_o(x)$ | S cm<sup>-1</sup> | $x$-dependent conductivity density of extracellular space in PRL | 1b, 3, 13a,b | Fig. 2 |
| $g_{i,k}(x)$ | S cm | Internal conductance of rod $k$ at position $x$; ($\text{M}\Omega \mu\text{m}^{-1}$ in figures) | 10 | Fig. 2 |
| $c_{m,k}(x)$ | F cm<sup>-1</sup> | Capacitance density function of rod $k$ of the ensemble | 22, 23 | Fig. 2 |
| $f_{\text{Kv2.1}}(x)$ | # | Distribution of Kv2.1 channel conductance over RIS; $\int_{x_{\text{OS/IS}}}^{x_{\text{ELM}}} f_{\text{Kv2.1}}(x) dx = 1$ | 5 | |
| $f_{\text{K, Leak}, k}(x)$ | # | Distribution of “leak” K<sup>+</sup> channel conductance; $\int_{x_{\text{OS/IS}}}^{x_{\text{ELM}}} f_{\text{K, Leak}, k}(x) dx = 1$ | Cf Eq 5 | |
| $f_{\text{HCN1}, k}(x)$ | # | Distribution of “leak” K<sup>+</sup> channel conductance; $\int_{x_{\text{OS/IS}}}^{x_{\text{ELM}}} f_{\text{HCN1}, k}(x) dx = 1$ | Cf Eq 5 | |

#### Structural properties

The ensemble comprised 11 model rods with identical outer and inner segments, and cell bodies uniformly spaced over the ONL (Fig. 2A-C; Table 1). The thickness of the C57Bl6/J mouse rod layer (*L*) was taken from histology and *in vivo* optical coherence tomography (Fig. 1). The axial resistance (*R*_i_(*x*), Fig. 2B) and capacitance (*C*_m_(*x*), Fig. 2C) profiles of the rods were derived from mouse rod EM data (Carter-Dawson and LaVail, 1979) and the assumptions that the cytosol has a resistivity of 115 Ω cm (Fry et al., 2012) and the plasma membrane a specific capacitance of 10 fF μm^−2^ (1 μF cm^−2^). The axial distributions of external conductivity and of the ionic mechanisms underlying *I*_m,*k*_ (Eq 3) were based on mouse retinal histology and recent literature (Fortenbach *et al*, 2021), as elaborated in the Results (Figs. 3, 4). The dimensions of the rods, capacitance distributions and axial resistance profiles are described in Figs. 2, 3 and Table 1.

**Figure 3.**
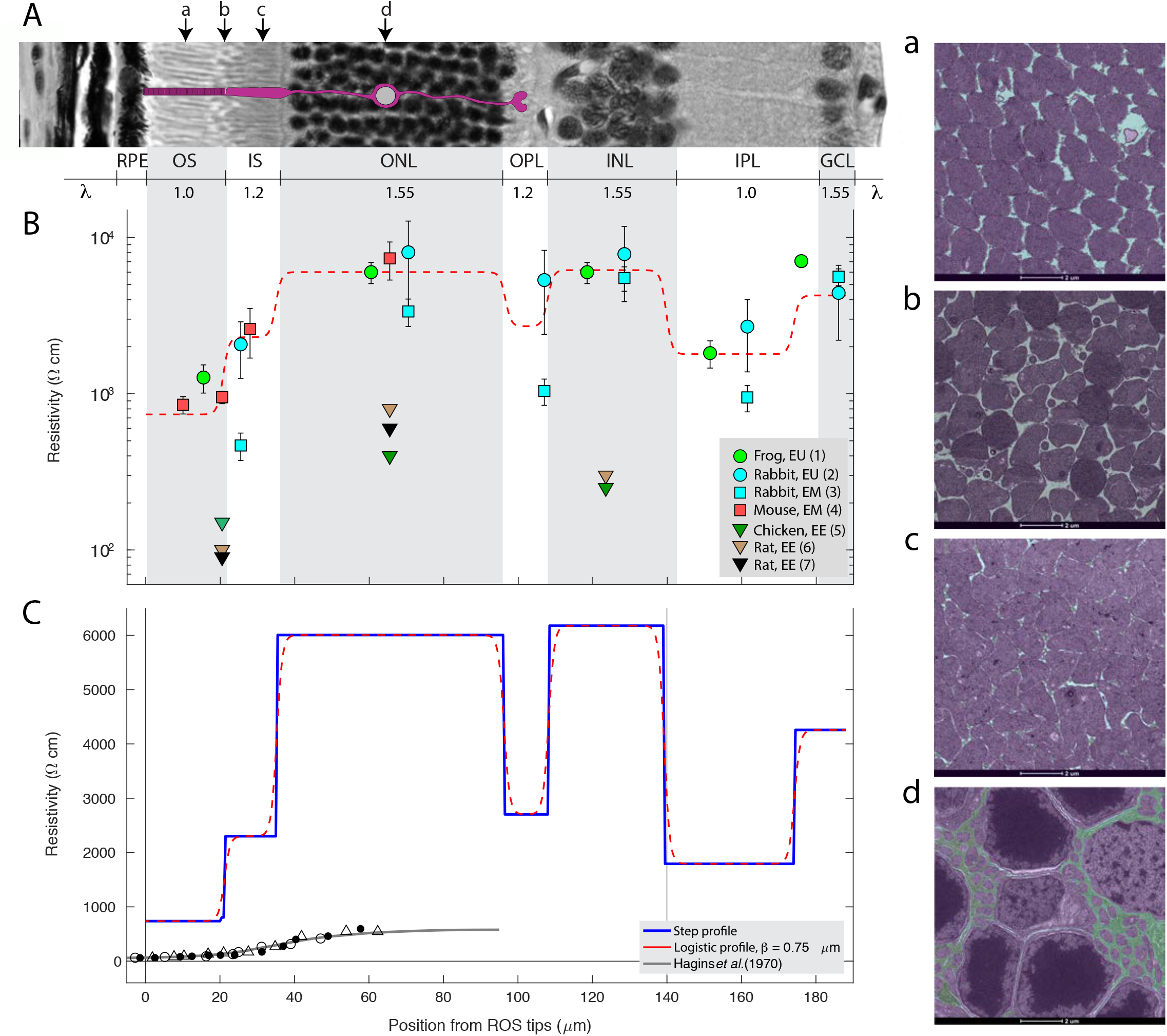
Axial resistivity profile of the mouse retina. **A**. Portion of the micrograph of Fig. 1A turn on its side to correspond to the spatial coordinate system; the 4 downward arrows (a, b, c, d) identify the depths of the electron micrograph sections at right. **B**. Resistivity profiles of rodent retina derived from several studies with the measurements derived from electrical measurements in Ussing chambers (“EU”), from electron microscopy (“EM”) (Eq. X) and from other electrical methods (EE), plotted on a logarithmic scale. There is a large range, with studies employing larger electrodes and electrodes reporting lower resistivities. **C**. Resistivity profile of the photoreceptor layer on a linear ordinate scale of a rat retinal slice preparation (symbols at bottom from 3 preparations) (Hagins et al., 1970) compared with that derived from the EM measurements in panel B. The blue line describes stepwise changes in resistivity between layers, while the red dashed curve (repeated in B, C) shows a version convolved with a logistic spatial filter.

**Figure 4.**
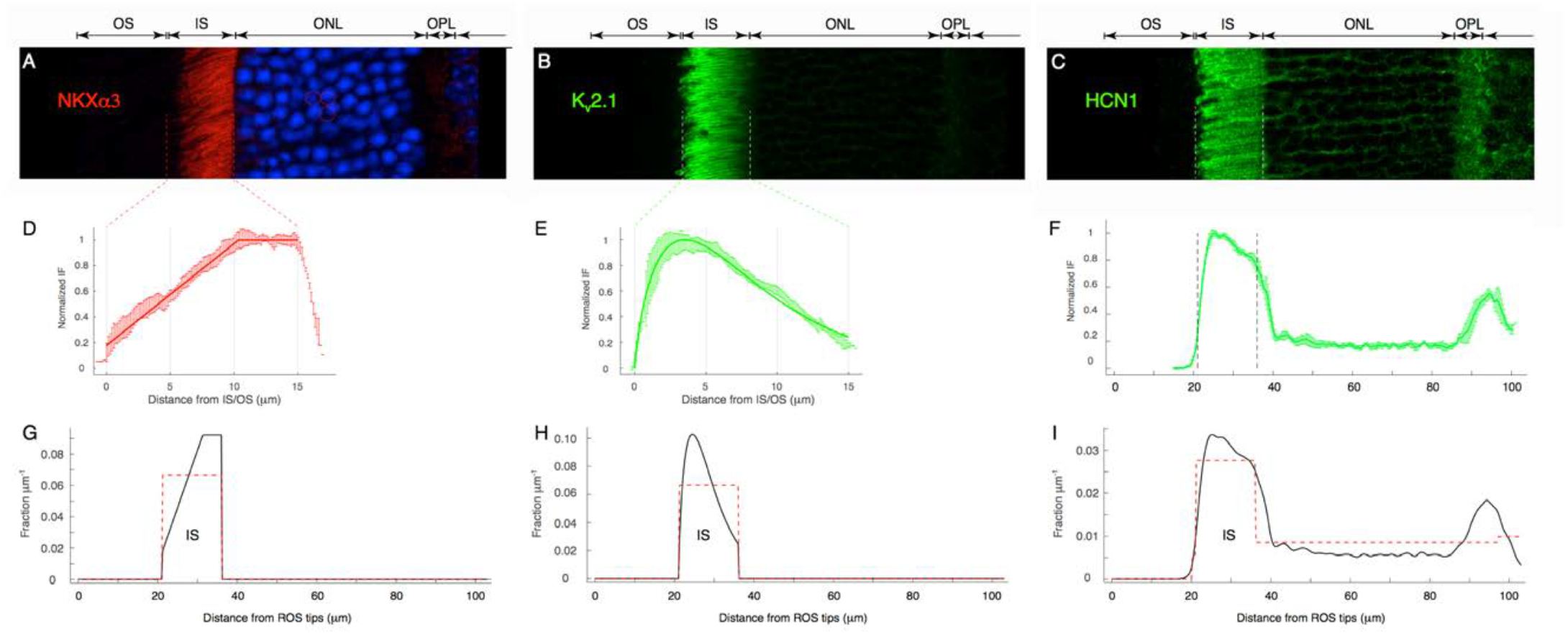
Spatial distributions of key non-outer segment ionic mechanisms contributing to the dark current and photoresponse of rods derived from immunohistochemistry. **A-C**. Confocal images of retinal sections immunostained for the NKX α3 rod-specific subunit (A), for K_v_2.1 (B) and HCN1 (C). The NKX and K_v_2.1 are expressed exclusively in the inner segment, while HCN1 is distributed throughout the non-outer segment region of the photoreceptor layer. **D-E**. Normalized immunofluorescence (IF) profiles of NKX and K_v_2.1 derived from digitally excised individual inner segments from sections (colored error bars are ± 2 SEM) (Spatially averaged IF from the IS layer gave similar results, but was further blurred on the left-hand side due to the variation in the position of the OS/IS.). Immunofluorescence (IF) of NKX and K_v_2.1 over the inner segments is non-uniform, and described by smooth curves. F. HCN1 immunofluorescence necessarily reflects averaging over different multiple rods. **G-H**. Density distributions over the entire rod layer used in the ensemble model derived from the profiles for NKX (G, black curve), and K_v_2.1(H, black curve). Uniform distributions (red dashed lines) are provided for comparison. I. The average IF profile of HCN1 over the photoreceptor layer is given by black curve; the dashed curves plot distributions used to assess whether the IHC-determined distributions are material to the behavior of the ensemble model.

#### Ionic mechanisms

The dynamical “forcing function” of Eq 3 comprises photo-transduction-driven changes in the CNG channel currents of the rod outer segments, and the consequent voltage-dependent changes in capacitive, ion channel and electrogenic exchange currents, which together comprise the full complement of sources and sinks of the ensemble. To make these latter explicit, the local membrane current density *I*_m, *k*_ (*x,t*) (A cm^−1^) of the *k*^th^ rod is expanded as follows:

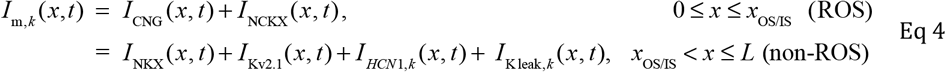

For clarity, in Eq 4 the domain of the spatial coordinate has been subdivided into the rod outer segment (ROS) region, and the rest of the rod layer (“non-ROS”). The CNG channel, NCKX currents and biochemistry of phototransduction machinery, which are confined to the ROS, were assumed to be identical for all the rods, as described in Gross et al (2012a, b) and (Fortenbach et al., 2021). Suppression of the rod index (*k*) for a particular ionic mechanism (e.g. the CNG channel current) indicates that all rods of the ensemble were assumed identical with respect to axial distribution and other properties of the mechanism. For the non-ROS region specification of the spatial distribution and membrane-potential dependence of the ionic mechanisms is required: e.g., the NKX transporter and K_v_2.1 channels are present only in the rod inner segment (RIS), while HCN1 channels are distributed throughout the rod outside the outer segment (Fortenbach et al., 2021) (Fig. 4) To allow some leeway in assessing the effect of different spatial distributions of the ionic mechanisms we created axial expression profiles – i.e, density functions whose integral over the spatial coordinate is unity. Thus, for example, the expression profile of K_v_2.1 channels is a function *f*_Kv2.1_(*x*) (units: μm^−1^) that satisfies 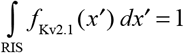 with the spatio-temporal K_v_2.1 current given by

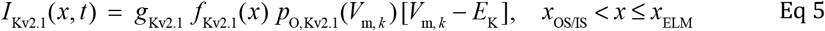

where *g*_Kv2.1_is the total (i.e., spatially integrated) K_v_2.1 conductance of the rod, *p*_O,Kv2.1_ is the instantaneous open probability at the local membrane potential *V*_m, *k*_ (*x,t*), derived from a 2-state Boltzmann dynamic model of Kv2.1 channels and [*V*_m, *k*_ − *E*_K_] the local electrochemical “driving force” for *I*_Kv2.1_(Beech and Barnes, 1989). A parallel expression applies for the HCN1 current density (Fortenbach et al., 2021). Equation 4 includes a small K^+^ “leak” current *I*_K leak, *k*_(*x,t*) whose existence was inferred from the analysis of currents of *K*_v_2.1^−/–^ rods (Fortenbach et al., 2021). The expression profiles of the electrogenic rod NKX, and of the K_v_2.1 and HCN1 channels will be described in a subsequent presentation of Fig. 4. The expression profiles of HCN1 and “K^+^ leak” channels varied from rod-to-rod in the ensemble due to the variation of the cell body location (Fig. 2A, B).

Specification of the ionic mechanisms requires explicit description of the voltage-dependence of their currents and activation and deactivation on membrane potential: for *K*_v_2.1 and HCN1 channels we used the 2-state Boltzmann models of (Fortenbach et al., 2021). The electrogenic Na^+^/K^+^ ATPase (NKX) was assumed to be the α3β2 isoform (McGrail and Sweadner, 1990; Schneider and Kraig, 1990; Wetzel et al., 1999). The dependency of NKX current on membrane potential was derived from the data of (Stanley et al., 2015), as previously described (Fortenbach et al., 2021). The NKX I-V curve was scaled so as to generate an outward current in the dark adapted rod (*V*_m, rest_ ~ −32 mV) corresponding to extrusion of all the Na^+^ that enters the ROS through the CNG channels and NCKX, assuming classical 3Na^+^ out: 2K^+^ stoichiometry for the NKX (Fortenbach et al., 2021).

Suppression of the CNG-activated current was assumed to follow “LP” kinetics (Lamb and Pugh, 1992; Pugh and Lamb, 1993)

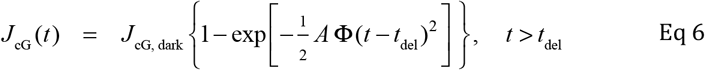

where *A* (s^−2^) is a constant characteristic of mouse rod phototransduction and *t*_del_ (s) a fixed, brief delay (~ 3 ms). The NCKX current was assumed to decay exponentially with a time constant of 80 ms after a saturating flash. As the CNG channels and the NCKX of rods have very weak dependence on membrane potential in the normal range of rod membrane potentials (Fortenbach et al., 2021), this description of the CNG and NCKX is appealing for its simplicity and established applicability: thus, here we are only concerned with the consequences for relatively brief times (< 50 ms) of the full closure of CNG channels in reponse to strong flashes.

Subject to the specifications just provided, we sought to solve Eq 3 for the axial- and time-dependent membrane potentials *V*_m, *k*_ (*x,t*) *k =* 1, …, *N*_R_, and for the external axial potential *V*_o_ (*x,t*) in the dark steady-state, and during the ensemble photoresponse to brief light stimulations of widely varied strength, i.e., for phototransduction-driven suppression of the rod CNG current for strong flash intensities. As next described, it was straight-forward to generate essentially continuous solutions of Eq 3 for the dark steady state. To obtain phototransduction-driven responses, however, it was found computationally necessary to “discretize” the spatial variable *x* into a limited series of positions, and to employ some additional mathematical tools, as described below.

### Steady-state solution for the dark adapted rod ensemble

In addition to ∂*V*_m, *k*_ / ∂*t* = 0, two general constraints must be satisfied by the dark steady state of Eq 3: (*i*) electrical balance of inward and outward current for each rod in the ensemble; (*ii*) homeostasis of the principal permeant cation species Na^+^, Ca^2+^ and K^+^. Formally, electrical balance means that the integral of the currents of the ion channels and electrogenic exchangers over the model rod membrane surface must be zero. Ion species homeostasis means that the net transmembrane flux of each permeant ion must be zero. Applying Eq 3 to test constraint (*i*) in effect reduces to performing two successive integrations over the spatial coordinate. First, the extracellular radial current density *J*_o_ is obtained from the average of the spatial integrals of the membrane current densities,

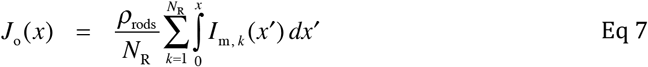

where *x* = 0 corresponds to the tips of the ROS, and *J*_o_ (0) = 0 (cf. Table 2 for units). The extracellular potential *V*_o_(*x*) is then obtained by a second spatial integration:

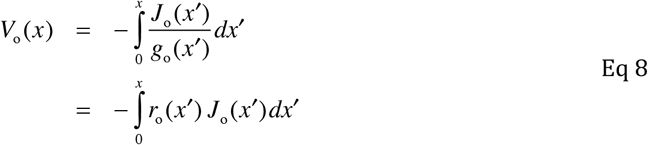

where *V*_o_ (0) = 0, and the second line serves to define the extracellular resistivity profile, *r*_o_(*x*) (units: Ω cm). Application of Eqs 7, 8 requires specification of the local ionic current densities *I*_m, *k*_ (*x*) and the membrane potential distributions *V*_m, *k*_ (*x*) of all rods of the ensemble. To address this requirement, it is useful to define the limb of the circulating dark current that flows internally in rods of type *k* of the ensemble (cf Table 2):

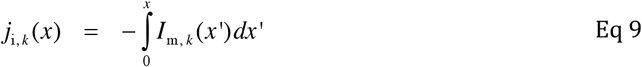

The internal current *j*_i, *k*_ (*x*) (units: A) is everywhere flowing in the positive direction by the choice of the spatial coordinate and the location of the reference potential at the rod tips, *x* = 0, and given the upper limit of the integral to be the total length of the rod layer, *L*, the integral must be zero by dint of the EOC. The corresponding internal axial potential of the *k*^th^ type rod is given by

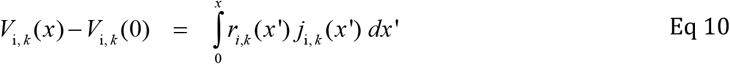

where *r*_i, *k*_ (*x*) (unit: Ω cm^−1^) is the internal resistance profile of the rod (Fig. 2B), and *V*_i, *k*_ (*x*) − *V*_i, *k*_ (0) is the potential difference that would be measured between a micro-electrode inserted at the tip of the ROS (*x*=0) and second microelectrode inserted at some more anterior location *x*. Because the system reference potential of the ensemble is located in the external medium at the ROS tips, *V*_i, *k*_ (0) = *V*_m, *k*_ (0). The steady-state membrane potential distribution is thus completely determined by Eqs 7 – 9 once a value is assigned to *V*_m, *k*_ (0) since by definition

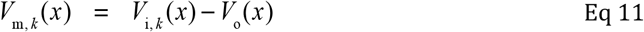

The value *V*_m, *k*_ (0) was determined by the constraint that the resting potential measured by point voltage-clamp at the rod cell body was –32 mV (Fortenbach et al., 2021) (cf. Fig. 5).

**Figure 5.**
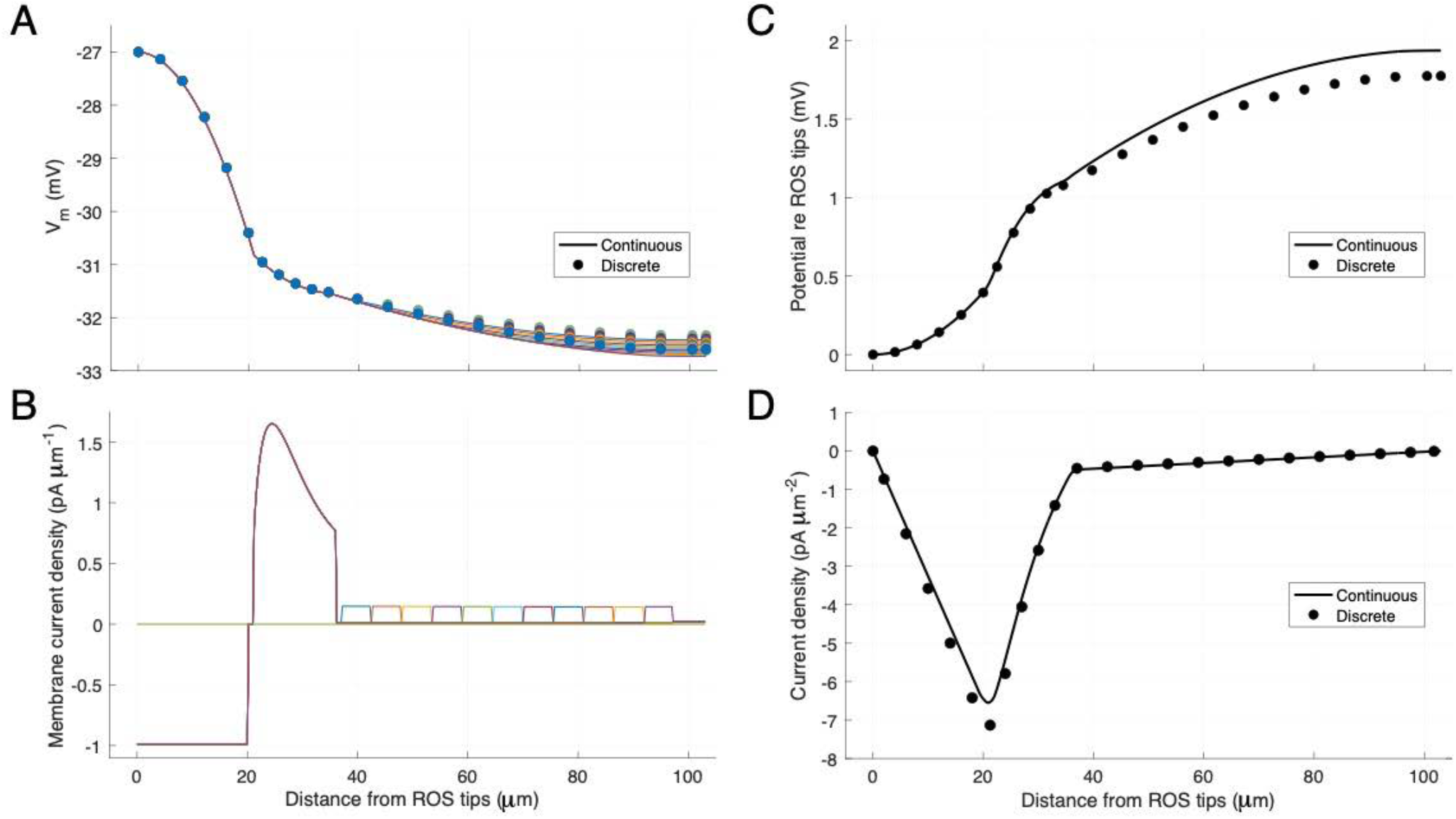
Dark steady-state distributions of the membrane potential and currents of the ensemble model. **A**. Membrane potential distributions of the 11 rods of the ensemble: smooth curves of different colors plot the “continuous” version of the model (d*x* = 0.25 μm), while symbols plot the results for the discretized, 24-node model. **B**. Membrane current density along the rod axis for the continuous version of the model. The uniform inward current of the outer segment integrates to −20 pA, and the non-uniform out-ward currents of the various rods each integrate to +20 pA so that the full spatial integrals for all rod are within 0.001% zero, i.e., obedience to the Equation of Continuity. **C**. The external axial potential of the ensemble for both continuous (curve) and discretized (symbols) model variants; deviation between continuous and discrete versions are a consequence of the discretization. **D**. Radial external current density of the ensemble for continuous and discrete model variants (1 pA μm^−2^ = 100 μA cm^−2^; cf Fig. S1). The sign of the current indicates the direction of flow: in the coordinate system adopted, negative means toward 0, the outer segment tips.

The ionic currents (Eq 4) in the dark steady state were specified as follows. All rods were assumed to have the same ROS current, comprising uniformly distributed CNG channel and electrogenic NCKX currents that sum to a specific value. The relative amplitudes of the CNG channel and NCKX currents were determined by the assumption that 7% of the inward CNG current is carried by Ca^2+^, according to the steady-state calcium homeostasis relation (*f*_Ca_ *J*_CNG_)/2 = *I*_NCKX,tot_, with *f*_Ca_ = 0.07 (Fortenbach et al., 2021). Thus, for example, assuming the rod has a total ROS dark current of –20 pA, then *I*_CNG,tot_ = –19.3 pA and *I*_NCKX,tot_ = –0.7 pA. The total inward flux of Na^+^ into the outer segment was then determined as (*f*_Na_ *I*_CNG,tot_ + 4 *I*_NCKX,tot_)/*F*, where *F* is Faraday’s constant, and *f*_Na_ can be taken to be 1 – *f*_Ca_. (Because the NKX of the RIS membrane is the only known mechanism by which Na^+^ can exit, and the rod has no material Na^+^ influx other than that into the ROS, by dint of Na^+^ ion species homeostasis the electrogenic NKX with its 3Na^+^:2K^+^ stoichiometry necessarily generates 34% of the total outward dark current, i.e., 6.8 pA for a rod with a 20 pA total dark current. Of the remaining 13.2 pA outward current, it was assumed that ~85% (11.2 pA) was carried by K_v_2.1 channels, with the residual 15% (2 pA) “leak” current carried by an unidentified type of K^+^ channels (Fortenbach et al., 2021). Though present everywhere in the rod except the outer segment (Fig. 4C), HCN1 channels contribute negligibly to the dark current due to the relatively depolarized membrane potential and the proximity of their reversal potential to the resting membrane potential (Fortenbach et al., 2021). The hyperpolarization-activated HCN1 current plays an important role, however, in shaping the rod photovoltage response (Demontis et al., 1999; Demontis et al., 2002; Seeliger et al., 2011) and so must be included in Eq 4. The total HCN1 conductance was set close to the value (1.45 nS) determined from characterization of the I/V curve of the dark-adapted rod (Fortenbach et al., 2021).

The dominant sources of the resting outward dark current of mouse rods, the NKX exchanger (α3β2 isoform) and *K*_v_2.1 channels, are expressed exclusively in the inner segments (Fortenbach et al., 2021) (Fig. 4). As NKX and *K*_v_2.1 expression appeared to be non-uniform in immunofluorescence imaging, we quantified the latter to derive spatial density functions *f*_Kv2.1_ (*x*) and *f*_NKX_ (*x*) (Fig. 4D, E, E, H), and further used the immunofluorescence of HCN1 to rationalize the distribution *f*_HCN1, *k*_ (*x*) of the different rods of the ensemble (Fig. 4C, F, I). The hypothetical K^+^ “leak” channel was assumed to be uniformly distributed in the non-ROS, non-RIS plasma membrane.

The extracellular potential *V*_o_(*x*) and the membrane potential distributions *V*_m, *k*_ (*x*) for the dark steady state (Fig. 5A, C), were determined by an iterative search process (“Numerical Methods”): these potentials were effectively continuous: thus, while an axial step size Δ*x* = 0.5 μm was used, no differences in the derived potentials were discernible when smaller step sizes were used in the iterative searches.

### Phototransduction-driven responses of the rod ensemble

#### Discretization of the spatial coordinate

To generate spatio-temporal photoresponses of the rod ensemble, we “discretized” the spatial coordinate into a grid of *N*_*L*_ positions, 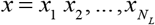 spanning the photoreceptor layer, i.e., *x*_1_=0 (ROS tips) and 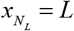 (inner retinal boundary of OPL and length of a rod). An electric cable circuit schematic of a single rod of type “*k*” (Fig. 2B-D) provides a helpful visual aid in the derivation of several key formulas, and in this framework each position *x*_j_ has a node for each rod. It bears emphasis that while a consistent 1-rod electrical cable circuit can be constructed, the *N*_R_-rod ensemble cannot be formulated as an electrical circuit. This follows because the interstitial current density that determines the extracellular voltage cannot be treated as the sum of the membrane currents of the rods of the ensemble, but rather is the sum of those currents weighted by the factor *ρ*_rods_ / *N*_R_ (Eq 6; as far as we are aware such a weighting has no parallel in electrical circuit theory).

We first insured fidelity of the discretized model with the effectively continuous model in the dark steady state (Fig. 5A, C). The discretized version of Eq 3 was then recast as a family of ordinary differential equations (ODEs), one for each of the *N*_*L*_ axial positions of each of the *N*_R_ rods comprising the ensemble, and the resultant set of ODEs integrated numerically as described further below. (In principle, discretization could be done with arbitrary *x*-resolution, but in practice spatial resolution was limited by computational capacity and speed.) The following sections provide the mathematical underpinnings of the discretization. The time variable is introduced to emphasize that the equations apply instantaneously.

#### Defining the rod ensemble model on a discrete spatial grid

The radial current density *J*_o_ (*x* _*j*_, *t*) (unit: A cm^−2^) of the ensemble at depth location *x* _*j*_ and time *t* can be represented as the average of the radial current contributions from each of the rod types:

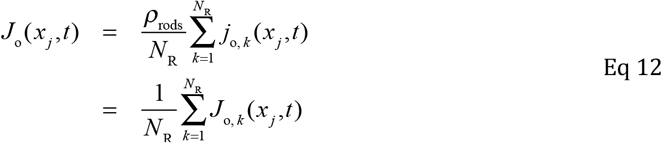

Here *J*_o, *k*_ (*x* _*j*_, *t*) ≡ *ρ*_rods_ *j*_o, *k*_ (*x* _*j*_, *t*) (A cm^−2^), i.e., is defined as the extracellular radial current density that would be generated were all rods of type *k*, and lower case *j*_o, *k*_ (*x* _*j*_, *t*) (unit: A) is used to indicate that the latter has different units. In the ensemble model, the component currents on the right-hand side of Eq 12 are cross-coupled through the dependence of the local membrane potentials of each rod on the common external potential *V*_o_ (*x* _*j*_, *t*) (Eq 1b). A theoretical potential *V*_o, *k*_ (*x* _*j*_, *t*) associated with the ECS current density of the *k*^th^ rod (cf Eq 1b) can be defined by

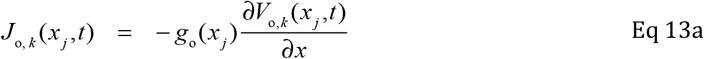

It is useful to rescale both sides of Eq 13a by dividing by the rod density ρ_rods_:

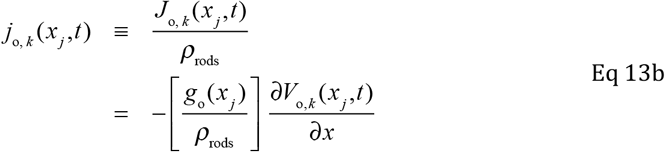

The units in Eq 13b of the reciprocal of the scaled conductivity *r*_o_ (*x*) = *ρ*_*rods*_ / *g*_o_ (*x*) (Ω cm^−1^) parameter are the same as those of the internal resistance of the rod, *r*_i, *k*_ (*x*) (Ω cm^−1^). To define the discretized model, these resistances were spatially integrated between the nodes, and the reciprocals of the internodal resistances used to define the conductance arrays *γ* _o_ (*x* _*j*_, *x* _*j*+1_), *γ* _i, *k*_ (*x* _*j*_, *x* _*j*+1_) (unit: S), *k* = 1, …, *N*_R_,; *j* = 1, …, *N*_L_. Instantaneously, the following Ohms Law relations must hold, starting with position *x*_1_ at the ROS tips:

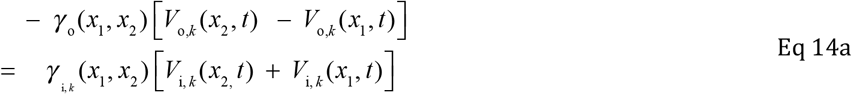

(with *V*_o, *k*_ (*x*_1_, *t*) ≡ 0 the ensemble reference/ground), while for an arbitrary axial position *x* _*j*_

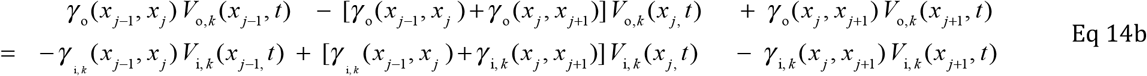

From the full set of these relations it follows that the column vector of external potentials 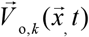 and that of internal potentials 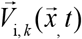 are related instantaneously to one another by a matrix equation,

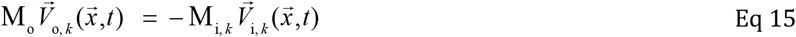

where the matrix

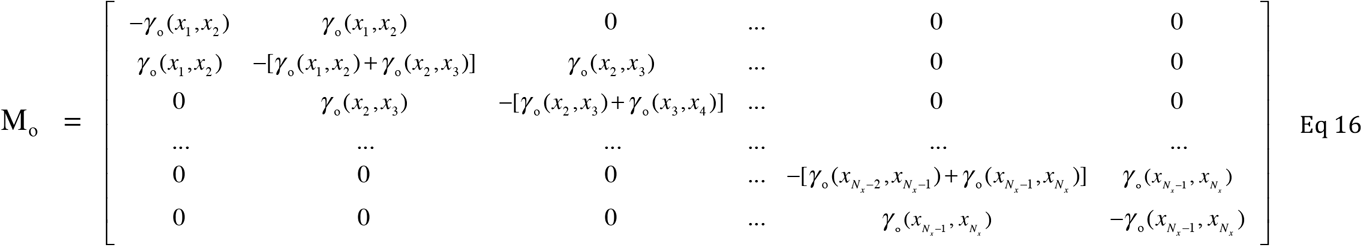

is not dependent on *k*. (For simplicity, we omit presenting an example of the matrices M_i, *k*_, but note that each has a symmetric structure like that of M_o_, with *γ* _o_ (*x* _*j*_, *x* _*j*+1_) replaced by *γ* _i, *k*_ (*x* _*j*_, *x* _*j*+1_)and that the first and last rows of each matrix are obtained from a relation similar to Eq 14a.) Averaging both sides of Eq 15 over the *N*_R_-rod ensemble, one obtains

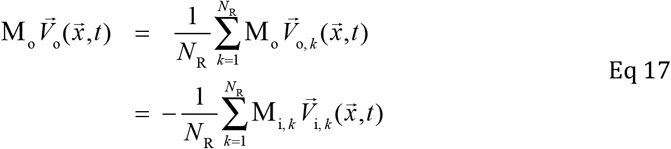

Equation 17 is a manifestation of the EOC in the discretized system: the external and ensemble-average internal “divergence” currents at each axial position *x*_j_ of the system must have equal magnitudes and opposite signs, and moreover, must equal the appropriately signed membrane current at that position. In the numerical implementation these relations were satisfied to within 0.2% at each *x*_j_ in the dark steady-state. (An explicit comparison of the continuous and discretized radial current distributions for the dark steady state is presented in Fig. 4C, and described in Results.)

The matrices M_o_ and M_i, *k*_ *k =* 1, …, *N*_R_ have the dimensions *N*_*L*_ × *N*_*L*_, and are singular with rank *N* − 1: consequently 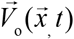 cannot be derived by multiplying both sides of Eq 17 by the inverse of M_o_. These matrices nonetheless have considerable utility in quantitative comparisons of the continuous and discrete models in the dark steady state, and in other derivations. In particular, the elements of the matrices can be used to construct a set of linear transformations T_*k*_ (*k* = 1,…, *N*_R_) between 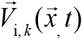 and 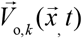 which provide a means for creating a system of ODEs that can be integrated to derive the ensemble’s photoresponses.

The construction of the transformation matrices T_*k*_ proceeds as follows. Letting m_o_ (*p, q*) represent the element of the matrix M_o_ with row and column indices *p, q* respectively, and similarly m_i, *k*_ (*p, q*) an arbitrary element of matrix M_i, *k*_, Eq 14a can be rearranged to give

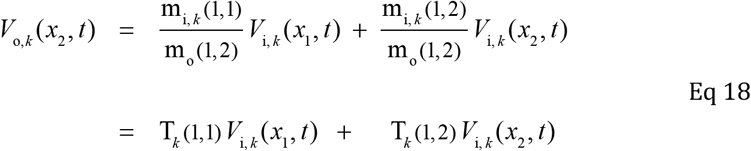

where the second line of Eq 18 serves to define the terms multiplying *V*_i, *k*_ (*x*_1_, *t*) and *V*_i, *k*_ (*x*_2_, *t*) respectively as the entries in columns 1 and 2 of row 1 of matrix T_*k*_; the remaining entries in row 1 of T_*k*_ are equal to zero. Next, rewriting Eq 14b in terms of the elements of matrices M_o_ and M_i, *k*_, and rearranging the terms to isolate *V*_o, *k*_ (*x*_3_, *t*) on the left hand side, one obtains

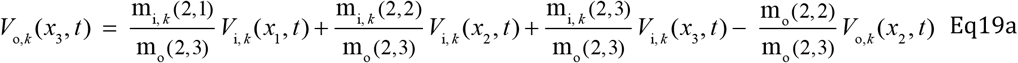

The term *V*_o, *k*_ (*x*_2_, *t*) is eliminated Eq 19a by substitution from Eq 18. Making this substitution, and collecting the terms that multiply the same internal potential variables *V*_i, *k*_ (*x* _*j*_, *t*), one obtains

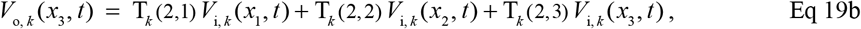

where

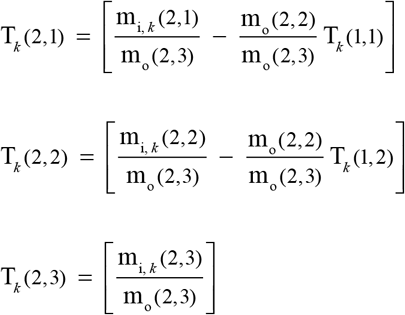

In this iterative fashion one obtains the entries for row 2, columns 1-3 of the transformation matrix, with the remaining row entries of zero. Applying Eq 19a recursively one arrives at a formulation for a physically dimensionless *N*_*L*−1_ × *N*_*L*_ matrix T_*k*_?that linearly transforms the *N*_*L*_ × 1 column vector 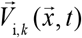 of instantaneous internal potentials of the *k*^th^ rod cell type into an (*N*_*L*_ − 1) × 1 column vector of external potentials, i.e.,

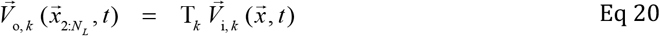

where 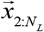 is used to indicate that the first element of the vector 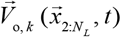 is the potential at the positions *x*_2_ to 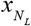, again keeping in mind that *V*_o, *k*_ (*x*_1_, *t*) ≡ 0, i.e. the external potential at the rod outer segment tips (*x*_1_) is defined as the system ground.

The utility of Eq 20 can be appreciated by its application to the full rod ensemble. Thus, the external potential of the ensemble of *N*_R_ rods is the average of the transformed internal potentials:

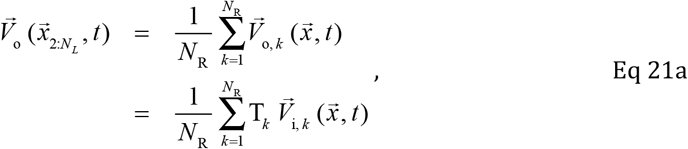

or, expressed for a single axial depth position *x* _*j*_, 2 ≤ *j* ≤ *N* _*L*_,

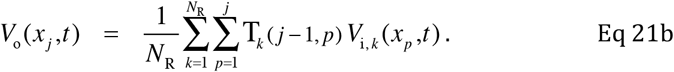

The restriction of the second summation in Eq 21b to terms with indices *p* ≤ *j* serves to emphasize a key feature of the construction of the matrices *T*_*k*_ (Eqs 18-19); the summation can be extended to *N*_L_ with no consequence, however, since *T*_*k*_ (*j* − 1, *p*) = 0, *p* > *j*.)

Eq 21 applies instantaneously at all times, including of course in the dark steady state. The prediction for the discretized ensemble of Eq 21 for the dark steady state is compared with the continuous prediction of the EOC in Fig. 5C. Reasonable agreement between the continous and discrete values confirms that discretization of the system has been implemented correctly, and that the spatial resolution of the discrete system is adequate. The interdependence of the different rods’ membrane potentials on *V*_o_(*x, t*) (which in turn depends on voltage-dependent membrane currents), makes it difficult to formulate a system of differential equations in terms of the rods’ membrane potentials. Another important consequence of Eq 21 is that it suggests that a dynamical (i.e., phototransduction-driven) discretized system can be formulated entirely in terms of the internal potentials, *V*_i, *k*_ (*x* _*j*_, *t*). Specifically, one can differentiate both sides of Eq 21b with respect to time and thereby express the time derivatives of the external potential in terms of the time derivatives of the internal potentials. This reveals that in principle the nodal potentials *V*_o_ (*x* _*j*_, *t*) and their derivatives can be formally replaced in the system of differential equations.

To see how a system of ODEs for the discretized *N*_R_-rod ensemble with *V*_o_ (*x* _*j*_, *t*) eliminated can be constructed, consider first the following partial differential equation:

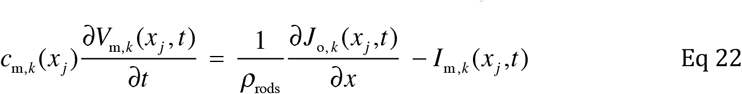

This equation separates out the contribution of type *k* rods to Eq 3, and rearranges the terms so that the time derivative of the local membrane potential is on the left-hand side. In this form the local capacitive current density (left-hand side of Eq 22) is seen to be equal to the difference between the properly (1 / *ρ*_rods_) scaled divergence (EOC) component and the ionic current density at depth position *x* _*j*_. The discretized version of Eq 22 is then written as

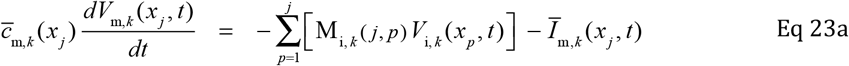

where the bars over the capacitance and ionic membrane current terms indicate that they represent the spatially integrated contributions of the internodal region bracketing location *x* _*j*_ (cf. Fig. 2), with the result that both sides of Eq 23a have the units of A. The summation term in Eq 23a corresponds to the “divergence” term of Eq 22. Given the identity*V*_m, *k*_ (*x,t*) = *V*_i, *k*_ (*x,t*) − *V*_o_ (*x,t*), Eq 23a can be rearranged as follows,

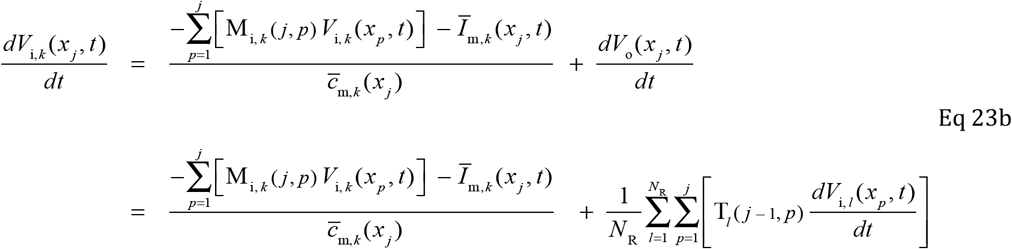

where the second line of Eq 23b is obtained from the first by differentiating both sides of Eq 21b with respect to time and substitution for *dV*_o_ (*x* _*j*_, *t*) / *dt*. In Eq 23b the variables *V*_o_ (*x* _*j*_, *t*) and their derivatives have been eliminated, creating a system of ODEs expressed completely in terms of the *N*_R_ × *N*_L_ variables *V*_i,*k*_ (*x* _*j*_, *t*), *k* = 1,…, *N*_R_, *j* = 1,…, *N* _*L*_. The second line of Eq 23b manifests cross-coupling between different types of rods, inasmuch as it shows how the internal potentials of all rod types in the ensemble affect *dV*_i, *k*_ (*x*_*j*_, *t*) / *dt*, the rate of change of the internal potential of the *k*^th^ rod at position *x* _*j*_. The final goal of the theoretical analysis is to establish a framework that enables the set of ordinary differential equations (ODEs) defined by (Eq 23b) to be integrated subject to their forcing by phototransduction-driven changes in CNG current.

#### Integration of the ODEs for the discretized N_R_-rod ensemble

Numerical integration of ODEs with predictor-correct methods such as Runga-Kutta requires the following criteria to be met: (*i*) initial values of all variables must be specified; (*ii*) a means of insuring that the spatial constaints between the variables are met at all times must be specified; (*iii*) given the values of the variables at any moment, the time-derivatives of those variables at that time point must be calculable, such that each variable’s derivative is isolated on the left hand side of its rate equation. The initialization of the internal potential variables *V*_i, *k*_ (*x*_*j*_, *t*) so as to satisfy all steady-state constraints of the dark adapted rods has been established (Fig 5), so criterion (*i*) is met. The spatial constraints interrelating the variables arise from the digitized divergence relation (Eq 11) by means of the terms 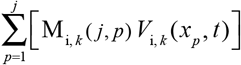 of Eq 22a, b, and so criterion (*ii*) is also met. A means of satisfying criterion (*iii*) was suggested by the fact that the identical expression for *dV*_o_ (*x*_*j*_, *t*) / *dt* is present in each of the set of ODEs (Eq 23b) for *dV*_i, *k*_ (*x* _*j*_, *t*) / *dt, k* = 1,…, *N*_R_. This insight leads to a matrix equation for the vector of rates of the internal potential changes at axial depth position *x* _*j*_ for the set of rods comprising the ensemble:

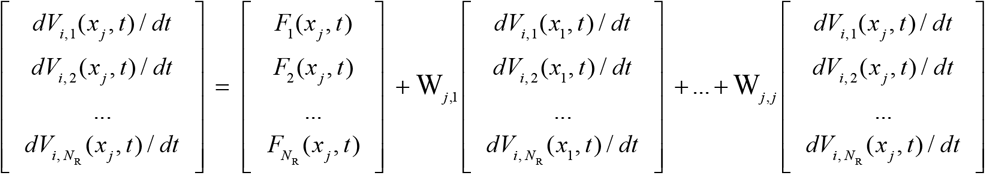

This relation can be succinctly expressed in vector format as

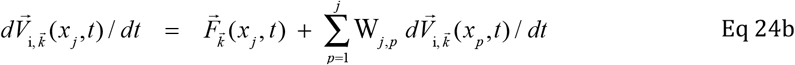

where 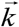 indicates that the vector 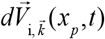 is “over” the set of *N*_R_ rod types, i.e., 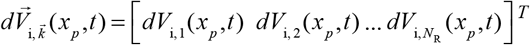 and the “nodal forcing functions” are

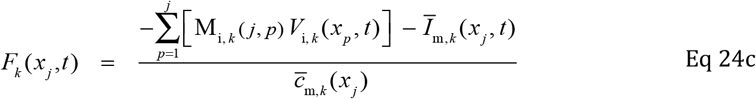

The matrices W_*j, p*_ (*j* = 1,…*N*_L_; *p* ≤ *j*) are *N*_R_ × *N*_R_ square matrices, with W_1,1_ empty (all zeros) because *dV*_o_ (*x*_1_, *t*) / *dt* = 0, and W_*p, j*_ determined from the transformations T_*k*_ (*k* = 1,…, *N*_R_) as follows:

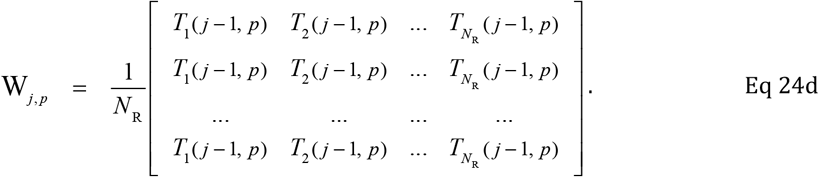

For any specific depth position *x* _*j*_ the vector 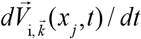 appears on both left- and right-hand sides of Eq 23b. However, it can be shifted entirely to the left side by subtracting the matrix W_*j, p*_ from the *N*_R_ × *N*_R_ identity matrix I :

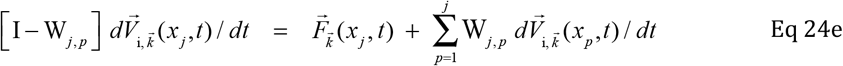

This construction creates a series of new matrices U_*j, p*_ ≡ [I − W_*j, p*_] that are non-singular and so invertible. Multiplying both sides of Eq 24e on the left by the inverse matrix U^−1^_*j, p*_ one obtains the following set of *N*_R_ × *N*_*L*_ ODEs:

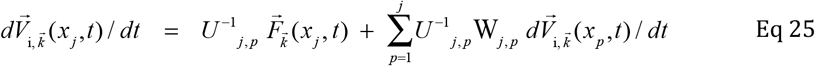

The critical feature of Eq 25 is that the *N*_R_ × 1 vector of derivatives of the internal potentials at depth location *x*_*j*_ of the rods specified by the left-hand side of the equations is only dependent on the right-hand side on functions of variables of the system at time *t*, and on the derivatives of the variables at spatial positions *x* ≤ *x* _*j*_, i.e., *p* ≤ *j* : thus, given *V*_i, *k*_ (*x* _*j*_, *t*), *k* = 1,…, *N*_R_; *j* = 1,…, *N* _*L*_, and the complete set of internal potentials at time *t*, all the factors on the right-hand side of Eq 25 can be calculated serially beginning at *x*_1_ = 0 and proceeding step-by-step to 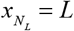. Consequently, the *N*_R_ × *N*_*L*_ equations specified by Eq 25 are seen to meet criterion (*iii*) stated at the beginning of this section for numerical integration with a Runge-Kutta or other predictor-corrector method.

To complete the specification of the dynamic model, rate equations for the open probabilities of the voltage-dependent *K*_v_2.1 and HCN1 channels at each spatial location must be included, and these probabilities used at each time step to compute the respective ionic currents (cf. Eq 5), and thus specify the terms 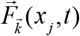. Notably, the various matrices involved in Eq 25 are purely “structural”, i.e., depend only on the external and internal conductivity structure of the ensemble of rods (Figs. 2,3; Table 1)and are not time-varying.

While the approach just described for solving the dynamic spatio-temporal photo-responses of rods *in vivo* is complicated, it has the virtues of being general, and applicable to any ensemble of electrically active cells with local lateral symmetry, i.e., collapsible to 1 spatial dimension. In particular, it resolves the knotty problem of the cross-coupling that occurs through the time- and space-varying external potential *V*_o_(*x, t*). Finally, with sufficient computational capacity, the discretization could be applied on such a fine grid that it would be effectively continuous. Thus, the method described provides a framework for similar applications of “source-to-tissue” CSD in other electrically active tissues.

## METHODS

All animal husbandry and procedures for animal handling during experimentation were as described in the preceding companion paper (Peinado Allina, 2025). Likewise, the methods of electrophysiological recording etc. were as described, as were the methods of light delivery and calibration. In the following we present methods that are either unique to this paper, or common to this paper and a subsequent companion paper.

### Electron microscopy and measurement of the retinal cross-sectional area occupied by the extracellular space

Mice were sacrificed by C0_2_ narcosis and perfused trans-cardially with 2% paraformaldehyde, 2% glutaraldehyde, and 0.05% calcium chloride in 50mM MOPS, pH7.4, as described elsewhere (Ding et al., 2015). Enucleated eyes were post-fixed in the same fixative overnight, after which the anterior segment removed and the eye cups diced into four 1mm x 1mm cubes. The tissue was then embedded in paraffin and aligned under the microtome to obtain slices cut perpendicularly to the longitudinal axis of photoreceptors. Starting at the outer nuclear layer, four 100 nm thick sections were collected on copper grids every 4 μm. Fifteen sets of slices were obtained spanning a total of ~60 μm, which was visually confirmed to correspond to half-way through the RPE (cf. Fig 3). Slices were stained with 4% uranyl acetate and 0.2% lead citrate.

Images were acquired with a FEI Talos L120C transmission-electron microscope. To avoid over-estimating the fraction of extracellular space at high magnification and under-estimating it a lower magnification, ROIs were imaged at three different magnifications (2600x, 5300x, 8500x), with 3 to 6 ROIs imaged at each retinal depth. The images were processed at full resolution manually in Adobe Illustrator to trace the extracellular space. Image masks were generated that assigned a value of 0 to every pixel corresponding to the extracellular space, and a value of 1 to all other pixels. The tabulated data were exported to Igor Pro to calculate the fraction of the imaged area occupied by the extracellular space.

### Pharmacological blocking of post-receptor currents by intravitreal injections

Intravitreal injections were performed under deep red light (>680 nm) to maintain dark adaptation. On the day of an experiment mice dark adapted overnight were anesthetized as described above and placed under a stereoscope on a regulated heating pad to regulate body temperature at 37°C. After inducing proptosis with mydriatics, methylcellulose was applied to the eye and an insulin needle was used to puncture the eye at the ora serrata. A gauge 33 metal tube was then inserted into the eye to deliver the solutions containing pharmacological agents. For all the experiments, a 4 μl volume was injected into the eye at a rate of 50 nl s^−1^ by means of a foot-triggered pump (UltraMicroPumpIII, World Precision Instruments). A second puncture was made at the top of the cornea to relieve intraocular pressure during the injection. As total volume of the mouse vitreous has been reported to be 4.4 ± 0.7 μl (Kaplan et al., 2010), a near-total exchange of the vitreal fluid was expected. After injection, animals were allowed recovery for 10 minutes in the dark before ERG recordings commenced. Recordings made after injections were used for analysis if the rod *a*-wave obtained in response to a flash producing ~ 4×10^5^ photoisomerizations per rod was maintained (Peinado Allina, 2025).

### Pharmacological agents and manipulations

DL-AP4, PDA and chloroquine were injected to blockers of photoreceptor synaptic transmission and Müller cell Kir7.1 channels in the photoreceptor layer. DL-AP4 sodium salt (Abcam Biochemicals), 2,3 cis-Piperidine dicarboxylic acid (Abcam Biochemicals), Chloroquine phosphate (Sigma-Aldrich), were all prepared with the same procedure: each drug was solubilized at 100 mM in ultrapure, molecular biology grade phosphate-buffered saline (PBS, Affymetrix/USBTM: 8 mM Na_3_PO_4_, 1.9 mM K_3_PO_4_, 2.7 mM KCl and 137 mM NaCl), and NaOH or HCl added to titrate the solutions to pH 7.3. Solutions were filtered with a 0.22 μm PES filter (Olympus plastics, Genesee Scientific Corporation), split into 50 μl aliquots, and stored at 4C. All solutions were prepared within a week of use.

Control experiments were performed to determine the minimum quantities of the drugs required to exhibit an adequate effect for the duration of a typical recording session (~60 minutes). An effective solution of AP4 and chloroquine suppressed the *b*-wave for a complete session. The effect of PDA (an iGluR blocker) could not be studied in dark adapted conditions. The effect of this drug was assessed based on the response to flashes delivered in the presence of a rod-saturating background of light (510 nm, 4.5×10^4^ photons μm^−2^ s^−1^) in experiments lasting for up to 60 minutes.

### Numerical methods

*Iterative determination of the dark steady-state of the N*_R_-*rod ensemble*. With the value of *I*_dark_ = *I*_CNG_ + *I*_NCKX_ specified as 20 pA, the total influx of Na^+^ was determined and thus the equal and opposite extrusion of Na^+^, fixing the outward electrogenic NKX current to 34% of the inward dark current (cf. (Fortenbach et al., 2021). Next the fraction of non-NKX current carried by K_v_2.1 channels was set to a value between 80% and 85% (in all results reported here, it was 85%), and the residual small outward current attributed to K^+^ “leak” channels. This resulted in a single conservation relation that needed to be satisfied for each of the rods of the ensemble (Eq 26):

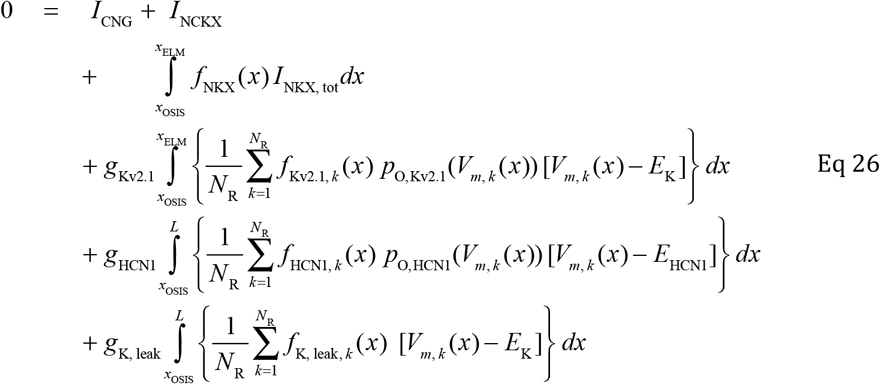

Here *I*_NKX,tot_ is the total outward electrogenic NKX current required to extrude the Na^+^ that enters the ROS through CNG channels and the NCKX, while the other terms are self-explanatory.

1. *Initialization*. The total inward current of the ROS, *J*_CNG_ + *J*_NCKX_, was set as described to the value −20 pA. A value for the membrane potential *V*_m, *k*_ (*x* = 0) at the ROS tips was selected (typically −27 mV); *V*_m, *k*_ (*x*) was thereby fully determined for the outer segments, 0 ≤ *x* ≤ *x*_OS/IS_ (the uniform current density in the ROS dictates that *V*_m, *k*_ (*x*) parabolically across the ROS; cf Fig. 5A). *V*_m, *k*_ (*x*) was initially assumed to be uniform at the value *V*_m, *k*_ (*x*_OS/IS_) for the rest of the rod, i.e., for *x*_OS/IS_ ≤ *x* ≤ *L*. The initial value for the total Kv2.1 conductance was chosen with the relation

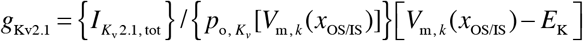

insuring that the total *K*_v_2.1 current was equal to that dictated by the fraction outward current selection. The residual outward current was assigned to the K^+^ leak channels by a similar relation, completing the initialization.
2. *Iterative searching*. In an iterative loop the right-hand side (RHS) of Eq 26 was repeatedly computed: if it exceeded 0, *g*_K, leak_ was decremented by a small amount Δ*g*_K, leak_, and vice versa if the integral was less than 0. In each iteration cycle the spatial distribution of the membrane potentials of each rod was calculated, the membrane-potential dependent open probabilities of the channels, etc., including small adjustments of *g*_Kv2.1_. This process was repeated until a sign reversal occurred. Upon sign reversal, the step size of Δ*g*_K, leak_ was reduced, and its sign changed. With this search process values of *g*_K, leak_ and *g*_Kv2.1_ was usually found within 20-30 iterative cycles that satisfied the EOC (charge balance) for all rods to less than 0.01% of the magnitude of the dark current. For the simulations reported in Figs. 5-9, the final values were *g*_Kv2.1_ = 0.264 nS and *g*_K, leak_ = 0.0469 nS for the continuous model, and the average fractional (relative to 20 pA) deviation over the 11 rods of the ensemble was 0.0004. A parallel iterative approach was used for obtaining the dark steady state of the discretized model, which provided the initial conditions for the rate equations of the model.

### Numerical solution of the photoresponses of the N_R_-rod ensemble

#### Spatial grid and system of rate equations

The *N*_L_ = 24 locations comprised 6 in the ROS layer, 5 in the RIS layer, 11 in the ONL corresponding to the centers of the cell bodies of each of the *N*_R_ = 11 rods of the ensemble), and 2 at the inner retinal boundary of the rod synaptic region, the OPL (Fig. 2). Thus, to describe the time-varying internal potentials *V*_*i,k*_ (*x* _*j*_, *t*), *k* = 1,…, *N*_R_; *l* = 1,…, *N*_L_ required *N*_L_ × *N*_R_ = 264 ordinary differential equations (ODEs), though this was effectively reduced by the assumption the ROS and RIS were identical for all rods of the ensemble. In addition 55 ODEs were required for the transition-rate equations for the 2-state Boltzmann Kv2.1 channel model in the RIS layer and 198 ODEs for the HCN1 2-state model over the “non-ROS” region (the 2-state Boltzmann model used are identical to those described in (Fortenbach et al., 2021).

#### Integration of the differential equations

With the initial conditions of all variables and parameters set by the steady-state analysis, the system of ODEs was integrated with the Matlab™ predictor-corrector script “ode23s”. The solution variables were used to generate the spatio-temporal membrane potentials of the 11 model rods of the ensemble, the varying open probabilities of the voltage-gated Kv2.1 and HCN1 channels, the membrane currents, etc. (cf. Figs. 7-9) for plotting and further analysis.

## RESULTS

### Estimation of the axial conductivity profile of the extracellular space of the mouse retina

In a transversally symmetric volume conductor such as the retina, the local conductivity *g*_o_(*x*) (units: S cm^−1^) is a scalar proportionality factor between the extracellular electric field 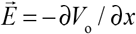 and the radial current density 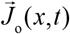 (Eq 1); resistivity is simply the reciprocal of conductivity, i.e., *r*_o_ = 1/ *g*_o_ In their classic investigation (Hagins et al., 1970a) directly measured *r*_o_ (*x*) as the resistance between a pair of electrodes spaced 10 μm apart and translated axially through a retinal slice. In comparison of published results from several species (Fig. 3B) we found the absolute magnitude of Hagins *et al*.’s measurements (Fig. 3C) to be questionable, and given the critical role conductivity plays in CSD analysis, we revisited the issue. Resistivity in a tissue volume conductor obeys the following relation (Gardner-Medwin, 1980; Mathias, 1983; Sykova and Nicholson, 2008):

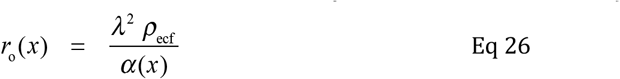

where α(*x*) (dimensionless) is the fraction of the tissue cross-section transverse to the *x*-coordinate occupied by the extracellular space, ρ_ecf_ (Ω cm) is the resistivity of the extracellular fluid, and λ is the dimensionless tortuosity factor (Gardner-Medwin, 1980; Mathias, 1983). We measured α in transverse EM images of the outer retina, derived resistivity with Eq 26, and compared our measurements with those of previous investigations (Fig. 3B). The studies of (Karwoski et al., 1985; Xu and Karwoski, 1994) are unique in demonstrating consistency between the total transretinal resistance measured in an Ussing chamber, and the summed layer resistances calculated from measured resistivity and layer thicknesses. These studies utilized used single thin electrodes (0.5 – 1 μm thick) to minimize disruption of the extracellular space in their measurement of resistivity. The estimates of resistivity from the EM studies (Reichenbach et al., 1988); Fig. 3B) and those of Karwoski *et al*. area also consistent with estimates of the tortuosity factor λ from studies of diffusion in cortical slices (Sykova and Nicholson, 2008).

To estimate the axial resistivity profile for the mouse retina (Fig. 3C), we combined the averaged mutually consistent estimates of layer resistivity (Fig. 3B) with the measured layer thicknesses of the mouse derived from OCT measurements (Figs. 1, 2; Table 1). Comparing these results with the measurements of rat retina (Hagins et al., 1970a) on a linear scale highlights a nearly 10-fold discrepancy (Fig. 3C). Notably, the three studies that reported relatively low resistivity (Fig. 3B, downward triangles) advanced multiple and relatively larger electrodes into the retina.

### Axial distributions of the Na^+^/K^+^ transporter (NKX), Kv2.1 and HCN1 channels

Calculation of the transretinal potential created by the rod dark current required specification of the axial distributions of the current sources and sinks of the rods, herein taken to be those of an 11-rod ensemble (Eqs 4, 5). Uniform distribution over the outer segment membrane of CNG (Cng1a, b) channels and the NCKX (Nckx1) electrogenic exchanger are well established and together comprise the inward dark current. The rod Na^+^/K^+^ ATPase (NKX, α3β2 isoform) and K_v_2.1 channels, which together account for about 85% of the balancing outward current in darkness, are completely localized in the inner segment (IS) membrane, while HCN1 channels are distributed throughout the non-OS membrane (Fortenbach *et al*., 2021). Quantitative analysis of immunofluorescence (IF) was used to derive axial distributions of the NKX, and of K_v_2.1 and HCN1 channels (Fig. 4). The IF of the NKX and of K_v_2.1 was distributed non-uniformly over the RIS (Figs. 4D, E), while HCN1 was distributed approximately in proportional to the calculated non-ROS membrane area (Fig. 4F). The measured IF distributions were used to create axial density function *f*_NKX_(*x*), *f*_Kv2.1_ (*x*) and *f*_HCN1, *k*_(*x*) (Fig. 4G, H, I) used in the model calculations. Mindful of the possibility of epitope artifacts, to assess the sensitivity of the model to the spatial distributions of the non-ROS ionic mechanisms, we also generated uniform density functions for NKX and K_v_2.1 (red dashed curves in Fig. 4G, H) and densities functions for HCN1 in the 11-rods proportional to the membrane surface area (Fig. 4I).

### Field potential of the rod circulating current: the dark steady-state

Given the spatial distributions of the ionic sources and sinks, the internal and membrane potential and current density profiles of the 11-rod ensemble were derived (Theory, Eqs 3-10), along with the extracellular current density and the extracellular potential for the dark steady-state (Fig. 5). Solutions were found iteratively for both the continuous ensemble model (unbroken curves) and for the discretized model (symbols). The external current density was close to that of (Hagins et al., 1970b) (Supplementary Fig. S1). These predictions are strongly constrained by a variety of published measurements of the voltage-dependence (and other relations) of the ionic mechanisms (cf. (Fortenbach et al., 2021)).

### Genetic and pharmacological isolation of rod-derived field potentials from the ERG

Electroretinogram (ERG) families were obtained from mice (*Gnat2*^−/−^) genetically engineered to lack the cone-specific G-protein and thus cone photoresponses (Ronning et al., 2018), and injected intraocularly with pharmacological blockers of synaptic transmission from photoreceptors to on-bipolar and off-bipolar cells (AP4 and PDA respectively) (Fig. 6). The effectiveness of the blockers can be appreciated by comparison of ERG response families with and without the injections (Fig. 6A, B): the explosive positive-going *b*-wave and accompanying oscillatory potentials are completely absent in the presence of the blocking agents (Fig. 6A, B). The *a*-wave, the initial corneal-negative response, was virtually identical in kinetics and amplitude with and without the blocking (Fig. 6C, D), revealing that the injection *per se* was not deleterious nor distorting. These results support the conclusion that post-synaptic signaling is negligible in the presence of the pharmacological blockers and the resultant ERGs arise from presynaptic rod activity. Nonetheless, the responses in the presence of the blocking agents exhibit a striking relaxation from their peak negative amplitude to a stable plateau. The negative-going transient and corresponding relaxation to the plateau has been called the “*a*-wave nose” and the pharmacological manipulations support the view that it originates in the rod photoresponses. The similarity of the *a*-wave nose to the HCN1-triggered depolarization following the hyperpolarization produced by strong light stimulation has often been noted, and it has been hypothesized that the *a*-wave relaxation and photovoltage nose may be causally linked (Demontis et al., 2002; Demontis et al., 2009; Turunen and Koskelainen, 2017). More recently, however, the *a*-wave nose has been attributed to the extracellular flow of capacitive current (Robson and Frishman, 2014) based on a cable model circuit analysis (cf. Fig. 2D). This latter model did not specify the molecular identities and voltage-dependencies of the underlying ionic mechanisms, and in particular did not include the HCN1 channels, which is needed to discriminate between the hypotheses. We thus sought to examine these hypotheses with the discretized ensemble model employing established rod ionic mechanisms (Fortenbach et al., 2021).

**Figure 6.**
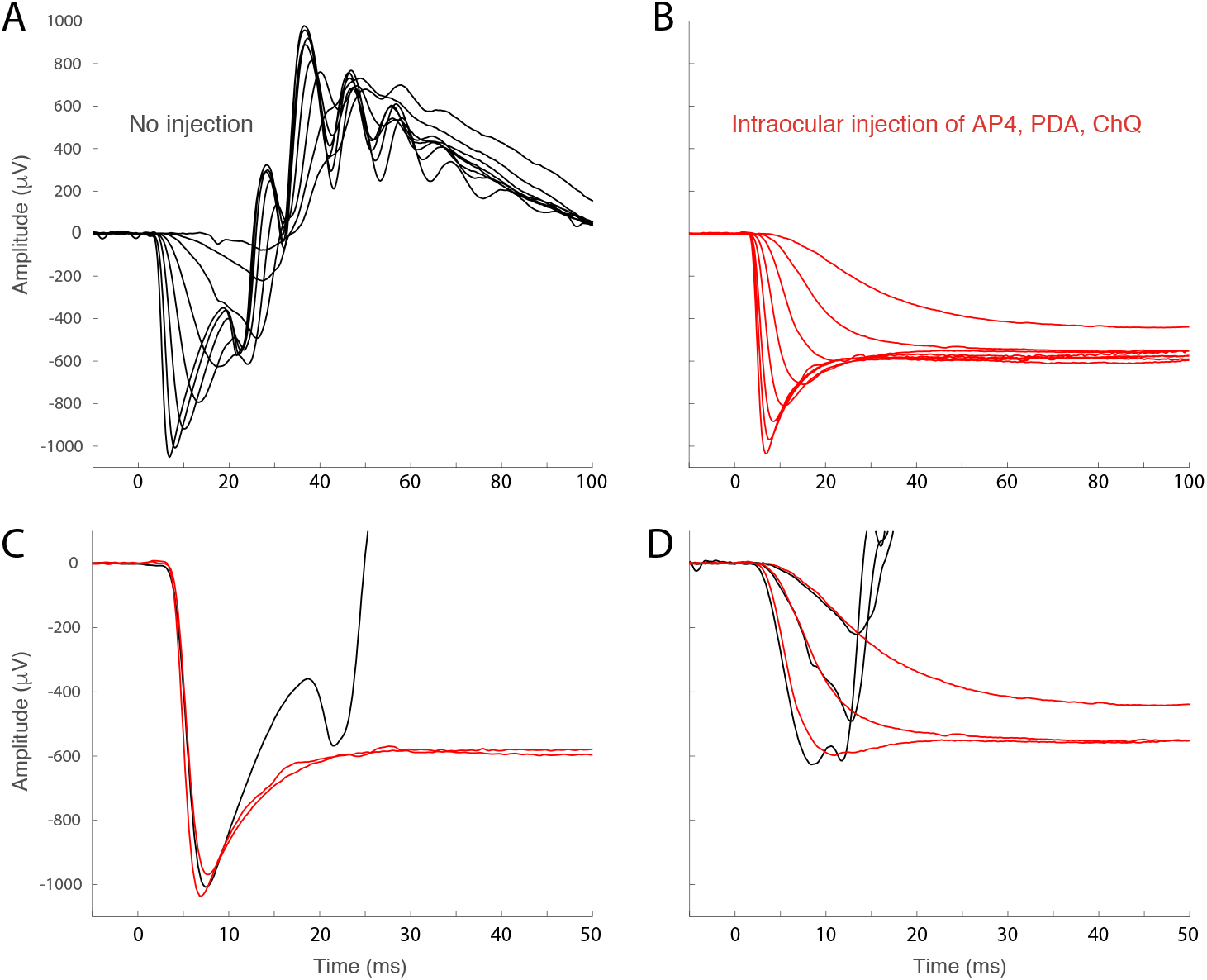
ERG response families of `Gnat2`^−/−^ mice with and without intraocular injection of the pharmacological agents to block post-synaptic mechanisms. **A**. ERG response family of `Gnat2`^−/−^ with no injection. **B**. ERG response family of mouse ~ 5 min after intraocular injection of AP4 (competitive inhibitor of glutamate binding to mGluR6), PDA (competitive inhibitor of bipolar iGluR channels) and Chq (competitive inhibitor of Kir7.1, expressed in Muller cells). The traces in A and B are plotted on the same ordinate scale. **C**. Comparison of the responses in A and B on an expanded time based to the most intense flashes – strong positive-going intrusion of post-synaptic responses is present by ~ 9 ms. **D**. Comparison of the responses in A and B to less intense flashes.

### Predictions of the rod a-wave by the ensemble model: membrane currents and extracellular potential changes

The *a*-wave response families of *Gnat2*^−/−^ mice with intraocular injections of synaptic transmission compared with predictions of the discretized 11-rod ensemble model (Fig. 7A). The theoretical traces reasonably recapitulate the key features of the *a*-wave family, including the saturated peak and plateau amplitude, and the kinetics initial phase of the *a*-wave traces over the 1400-fold intensity range. The predictions deviate from the traces, however, by undershooting the responses of all but those in response to the two most intense flashes. For the responses to the four lowest intensities, the deviations are expected, as photoactivated rhodopsin in mouse rods is deactivated with an apparent time constant less than 50 ms, and calcium-feedback activation of guanylate cyclase also occurs during the rod response rising phase (Burns and Pugh, 2009; Gross and Burns, 2010; Gross et al., 2012), while the “LP” kinetic model (Eq 6) of the light responses assumes no deactivation or calcium feedback. These omissions can be remedied, but for addressing the nature of the *a*-wave “nose” relaxation, it suffices to focus on the response to the most intense flash, estimated to produce 3.5 ×10^5^ photoisomerizations^−^ rod^−1^ and suppress all CNG current for more than 3 s *in vivo* (Peinado Allina et al., 2017, Fig. 2A). Various aspects of the model response to the strongest stimulus are provided, including the extracellular potential at various depths and times (Fig. 7B, C), the membrane potential of one of the rods of the ensemble (Fig. 7D), all the membrane currents of one of the rods of the ensemble (spatially integrated), and a dynamic test of the Equation of Continuity (Fig. 7F, dashed line). The latter (Fig. 7F) establishes that the predicted membrane currents of the outer segment and the rest of the rod sum to zero at every moment in time during the photoresponse. In addition to its closely replicating the measured saturated *a*-wave, two features of the theory trace (Fig. 7A & 7B, bright red trace) bear emphasis: first, the predicted trans-photoreceptor layer photovoltage (*V*_o_(*x*_L_, *t*), Fig. 7B, bright red) was scaled by a factor of 0.21 to match the data trace (cf. Discussion); second, the predicted trans-retinal photovoltage relaxes to a plateau below the level set by the absence of extracellular current in the photoreceptor layer (Fig 7B).

### Physiological mechanisms of the a-wave “nose”

Some insight into the way in which the extracellular current arising from different mechanisms of the rod contribute to the trans-retinal photovoltage can be obtained by considering the potentials that each would generate by itself (Fig. 8). This approach is possible because an instantaneous axial potential distribution associated with each component can be recovered from the model, and these potentials, like the underlying axial currents, must sum at each depth according to the Equation of Continuity. The baseline 22 mV potential of the CNG current (Fig. 8A) declines essentially as a step (Eq 6), with the other potential changes in effect revealing the “step response” of the model to its physiological step forcing function. The membrane potential (Fig 7D) declines in very nearly the same way for each rod (cf. Fig 7D), but its time course is determined not only by the “membrane “time constant” (~3 ms), but also by the kinetics of the voltage-dependent Kv2.1 channels’ (open to closed) and HCN1 (closed to open) channels’ state changes (Fortenbach et al., 2021, Fig. S2). The resting axial potentials of the two largest ionic non-outer segment components, Kv2.1 channels (Fig. 8C, −12 mV) and HCN1 channels (Fig. 8D, 0 mV), and the signs of their photopotentials are both positive. The photoresponse diminishes the amplitude of the Kv2.1 potential as these channels close in response to hyperpolarization, while the amplitude of the HCN1 potential increases as the latter channels open. Aside from the decline in the potential of the CNG current, the component of the photovoltage with the largest magnitude is the capacitive transient (Fig. 8E). The sum of all the component potentials (Fig. 8F) recapitulates the directly computed solution (Fig. 8B). An important confirmed result is that the transretinal potential is ~ − 1 mV at the innermost retinal level (*x*_L_= 103 μm) during when the saturated *a*-wave reaches its plateau.

**Figure 7.**
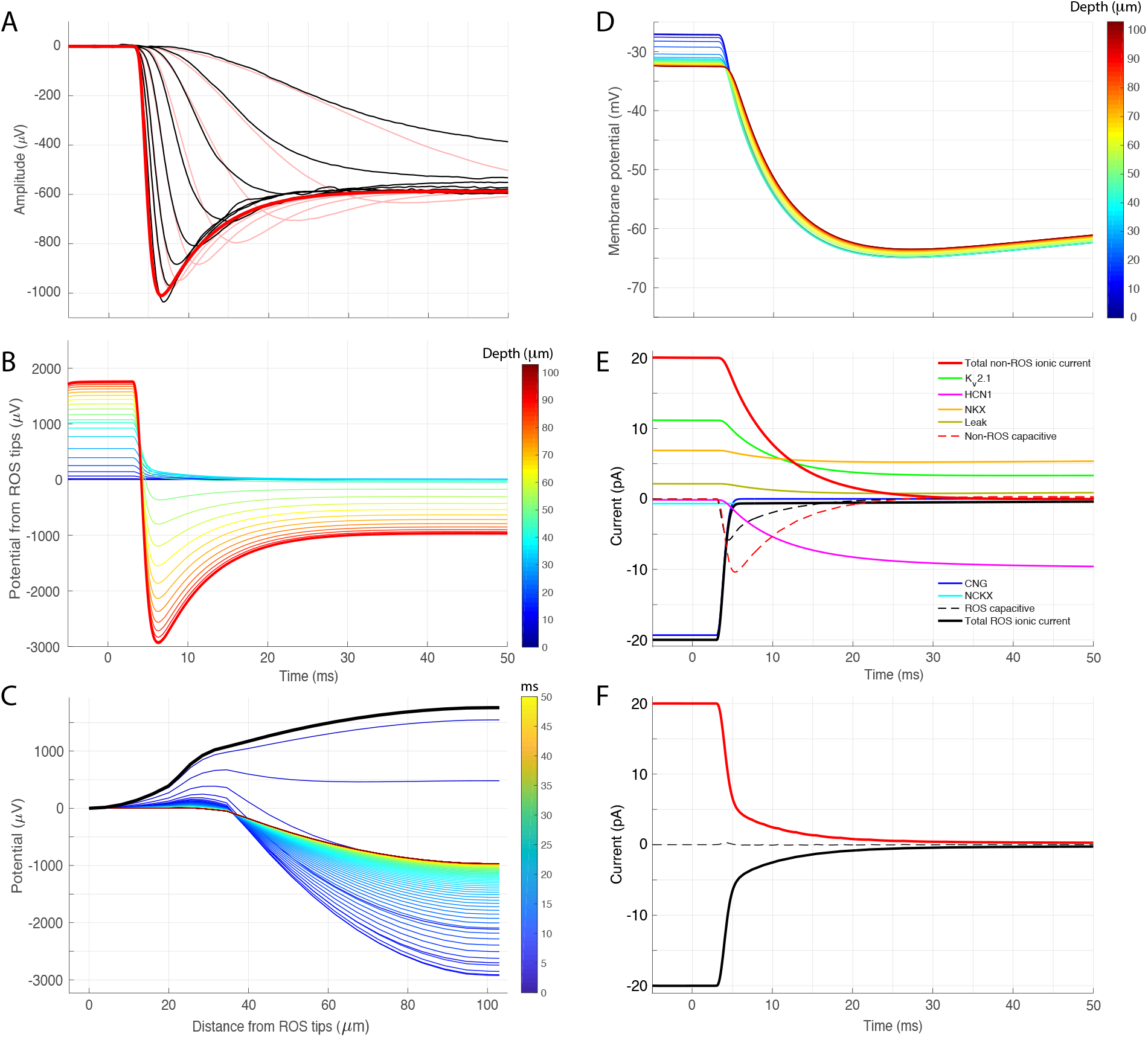
Rod ensemble model predictions of the ERG *a*-waves of `Gnat2`^−/−^ mice measured in the presence of blockers of synaptic transmission. **A**. Corneally recorded *a*-wave family (black traces) and the predictions of the 11-rod ensemble model (red traces; cf Theory). Flashes were estimated to generate between 250 and 3.5×10^5^ photoisomerizations per rod. **B**. Predicted photopotentials at different positions in the photoreceptor layer relative to the rod outer segment (ROS) tips for the response to the most intense flash in A (layer depth coded by trace color; *cf* colorbar at right). In panel A the red trace predicting the response to the brightest flash was created from the trans-photoreceptor layer potential (thickened red trace in B): for the prediction in A, the baseline of the latter trace was offset to zero and scaled by the factor 0.21. **C**. The results in B replotted as a function of the spatial coordinate with time as the parameter (color bar at right). The thickened black trace at the top replots the dark steady-state potential distribution arising from the dark current, and corresponds to the symbols in Figure 5C. (As the time increment of the numerical integration was 100 μs, the ODE solutions over the 55 ms epoch comprise 5501 time points; for clarity only 1 in 5 of the solution traces is shown, i.e., every 500 μs. **D**. Predicted membrane potential response of the model rod (#6) whose cell body is situated in the middle of the ONL. The traces are color coded for depth in the photoreceptor layer (colorbar at right), and vertical differences between the traces manifest spatial variation in membrane potential (*cf* Figs. S1, S2). **E**. Membrane currents of rod#6 predicted by the model for the response to the brightest flash. The color code for the different ionic mechanisms is identical to that used in Fig. 6 of (Fortenbach et al., 2021) where the voltage-dependence of the various ionic mechanisms is presented. (The ensemble model predicts the currents at each of the 24 axials “nodes” of the model, but for this plot they were summed “by ionic type” over the spatial coordinate.). Of particular note is that the K_v_2.1 (green) and (to a lesser extent) NKX (orange) and K^+^ “leak” currents begin declining when the CNG-channel current is suppressed (blue trace), and at the same time (but with a small time lag) the HCN1 current becomes inward and increases (magenta trace). The spatially summed capacitive currents of the outer segment and “non-outer segment” are also shown (dashed black and red traces respectively). **F**. Sum total ionic and capacitive membrane currents of the outer segment (black trace) and non-outer segment (red) for the predicted response of rod #6 of the ensemble to the brightest flash (these traces represent running sums of the respective currents in panel E). The sum of the inner and outer segment membrane currents (dashed line) is zero (except for a small deviation at the onset of the CNG channel response). The zero dynamic sum is physically required by the Equation of Continuity (EOC; Eq 2), and here reveals that the model formulation and coding are consistent with this requirement, while the effectively mirror image form of the currents of the outer segment and “non-outer segment” is a predictable consequence of the distribution of the current sources and sinks. Similar “dynamic conservation of charge” (obedience to the EOC) was obeyed for all rods and all flash strengths, and for each of the 24 nodes of the 11 rods.

**Figure 8.**
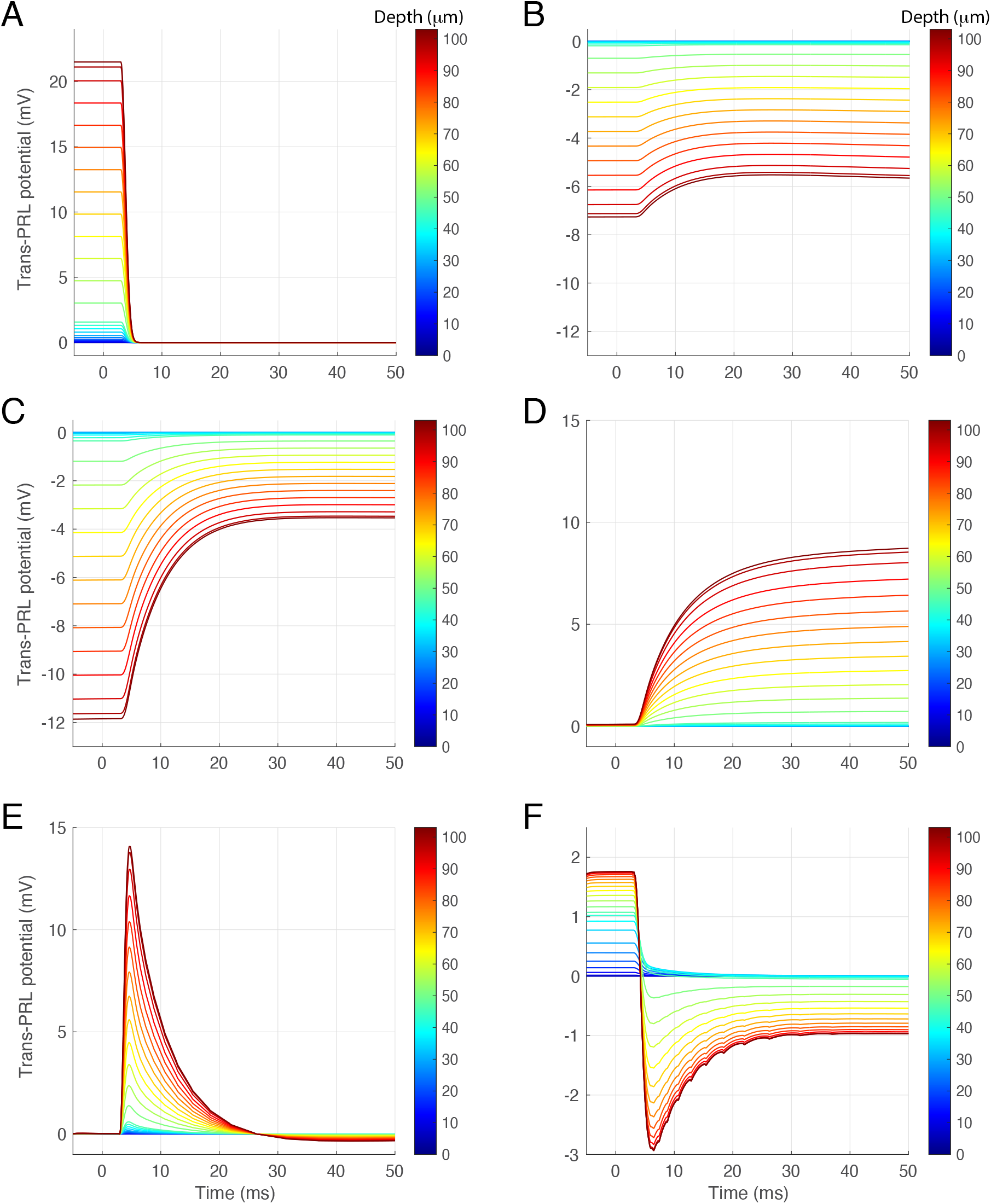
Trans-photoreceptor layer potentials of the molecular determinants of the saturated mouse rod *a*-wave (3.5×10^5^ photoisomerizations per rod; the colorbars indicate the depth in the photoreceptor layer of the traces.) **A**. Extracellular potential of the CNG channel current in response to a flash producing 3.5×10^5^ photoisomerizations per rod – cf. Eq 6). **B**. Extracellular potential of the NKX electrogenic current (cf. Fig. 4A, D, G). **C**. Extracellular potential of Kv2.1 channel current (cf. Fig. 4B, E, H). **D**. Extra-cellular potential of HCN1 channel current (cf. Fig. 4C, F, I). **E**. Extracellular photopotential of the capacity current. **F**. Depth varying photopotential generated by summing at each moment in time all component potential for each of the 24 depth locations spanning the photoreceptor layer (PRL). The red/black curve corresponding to 103 μm reproduces directly the trans-PRL photopotential directly computed with Eq 21b from the solution of the model rate equations (Fig. 7B, red curve): this correspondence is expected given the model’s dynamic obedience to the EOC, but serves here to demonstrate that the component potentials are correctly computed by integration of the individual underlying current components over the PRL conductivity distribution.

Further insight can be obtained by plotting together all the component photopotentials at the innermost retinal location, *x*_L_ (Fig. 9A) and deleting either the HCN1 contribution or the capacity current contribution (Fig. 9B). This reveals that both components make a material contribution to the composite trans-retinal photopotential that describes the saturated *a*-wave (Fig. 9C). For each moment in time extracellular currents corresponding to the component potentials of Fig. 8 can be derived by the numerical approximation of the volume current density relation (Ohm’s law for the digitized model; not shown). Doing so reveals that the extracellular capacity current flows toward the outer segment for ~ 20 ms (cf. Fig. 8E), and that once activated, the extracellular HCN1 current likewise flows toward the outer segment, while the K_v_2.1 component of extracellular current flows in the opposite direction, so that the currents of the two voltage-gated channels largely cancel, combining to form a net negative transretinal potential, the undershoot.

**Figure 9.**
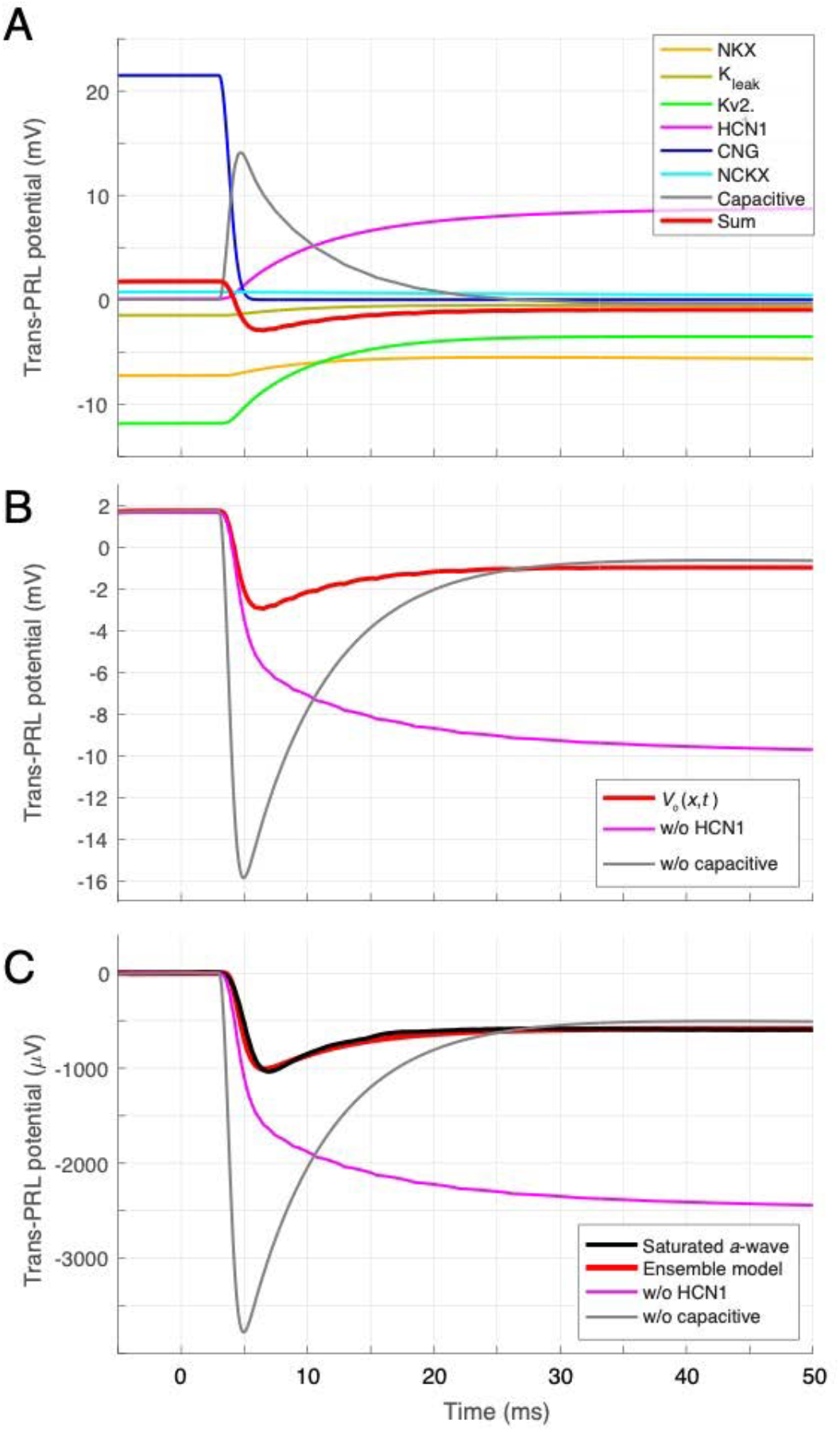
Assessment of the contributions of different rod mechanisms to the trans-retinal potential (TRP) underlying the saturated *a*-wave. **A**. Predicted photopotentials at the proximal PRL (103 μm) of all components contributing to the saturated *a*-wave. The color scheme is identical to that used in Fortenbach *et al*. (2021) to identify the same ionic mechanisms’ contributions to the I/V curve of the dark adapted rod, with the exception of the capacitive potential (gray curve). As the currents underlying these potentials summate in the extracellular space, so must their hypothetical potentials. Note in particular that the very fast and large negative-going CNG channel potential is largely neutralized by the positive-going capacitive transient. While similar cancellative behavior happens at the membrane (cf. Fig. 7E), the signs here provide the direction of the underlying current by dint of the Ohm’s law relation, *J*_o_ (*x,t*) = − *g*_o_ (*x*)[∂*V*_o_ (*x,t*) / ∂*x*], which express the principle that in volume conductors current flows “down the potential gradient” with local conductivity the proportionality factor. Thus, a positive potential indicates net current over the PRL flowing towards the RPE, whereas a negative potential indicates net current flowing toward the OPL (Fig. 1). **B**. Trans-PRL potentials revealing the effects of deleting the HCN1 component (magenta) or the capacitive component (gray). The red curve repeats the curve corresponding to the sum of all components (Fig. 7B, Fig. 8F, red/black curve). **C**. Comparison of the HCN1-deletion (magenta) and capacitive-deletion (gray) photopotentials and the sum of all components (red cuve) with the measured saturated *a*-wave (black trace). All curves have been scaled been offset to zero and rescaled by 0.21, the factor that brings the prediction into correspondence with the saturated *a*-wave.

## DISCUSSION

The goal of this investigation (Robson and Frishman, 2014)was to isolate the rod-driven field potentials of the mouse retina with genetic elimination of cone signaling (*Gnat2*^−/−^), pharmacological blockage of trans-synaptic signaling (AP4 and PDA), and to explain the amplitude and kinetics of the early ERG measured with corneal electrode (Figs. 6, 7) in terms of electrical (Fortenbach et al., 2021) and structural properties of the photoreceptor layer (Figs. 1-3). The 11-rod ensemble dynamic model created to achieve this goal encompasses the principal heterogeneity of rods – the location of the cell body at varying depths in the ONL (Figs 2, 3) – and mechanism-specific axial expression distributions for the ionic mechanisms extracted from high-resolution confocal immunofluorescence (Fig. 4). The model was able to recapitulate the principal features of the *a*-waves, including the light-dependence of the early phase, the amplitude of the plateau phase relative to the saturating negative peak (Fig. 7).

### The resistivity of the retina in vivo and the absolute scale of the corneal ERG a-wave

A definitive biophysical account should explain the absolute amplitude of the saturated *a*-wave responses, which with the air-gap electrode method (Peinado-Allina *et al*, 2025) is about 1000 μV (Fig. 6C, 7A). The theoretically predicted amplitude (Fig. 7B, red trace) is 4700 μV, while the observed amplitude at the cornea is ~1000 μV, implying an overall reduction factor of 0.21. The 4-shell model of the mouse eye predicts a reduction factor of 0.38 between a trans-photoreceptor layer potential and the corneal potential (Peinado-Allina *et al*, 2025), so that a residual 0.38/0.21 = 1.8-fold amplitude reduction remains unaccounted for. What factor(s) might account for this? The axial extracellular resistivity profile *r*_o_(*x*) is a critically important determinant of the magnitude of the field potential, and simulations with the profile derived from EM results (Fig. 2C) show that the residual discrepancy can be bridged by scaling *r*_o_(*x*) by ~ 0.6, suggesting that estimates of resistivity from EM are distorted by the fixation process. An alternative and not mutually exclusive explanation is that the effective resistance of the ora serrata region is lower by approximately a factor of 2 from that employed in the 4-shell model (Peinado-Allina *et al*, 2025). Another contributing factor may be the rod density: it could plausibly be 15% lower than the value adopted (3.5×10^5^ rods mm^−2^). Despite this absolute scaling discrepancy, the “forward CSD” modeling is consistent with the current density measured by Hagins *et al*. (1970) (Fig. S1), whose measurement depended on their measurement of the resistivity in the *ex vivo* preparation, not on that *in vivo*. Nonetheless, extraction of resistivity from EM and other data (Fig. 3), highlights the problem of inserting electrodes into the highly packed retinal parenchyma, and possibly into *ex vivo* retinal slices more generally.

### The mechanism of the a-wave “nose”: roles of capacitive and. HCN1 currents

When corneal ERGs are measured in response to increasingly intense flashes, the negative-going *a*-wave saturates, i.e., does not alter its amplitude or kinetics with further increasing flash strength (Fig. 6A)(Kessler et al., 2014; Robson and Frishman, 2014). With genetic (*Gnat2*^−/−^) elimination of contributions from cones, and pharmacological blocking of synaptic signaling (Fig. 6B), the isolated, saturated rod *a*-wave can be recorded (Fig. 6B, C). The saturated *a*-wave exhibits a “nose”, a striking relaxation from its peak magnitude to a stable plateau that is about 60% the peak magnitude. The strong light intensities that produces the saturated *a*-wave are known from single-cell and *in vivo* mouse rod recordings to very rapidly and completely close CNG channels for several seconds (Peinado Allina et al., 2017) – in effect producing a step-like “forcing function” driving all the molecular mechanisms underlying the rod photoresponse (cf. “CNG” curve in Fig. 7E), and causing the rod to hyperpolarize at its “maximum warp” (cf. Supplement, Fig. S2).

The *a*-wave “nose” has been hypothesized to arise from the activation of HCN1 channels (Demontis et al., 1999; Demontis et al., 2002; Turunen and Koskelainen, 2017) or from the extracellular flow of capacity current (Robson and Frishman, 2014). The rod ensemble model enables these hypotheses to be addressed by examination of the component trans-photoreceptor layer field potentials (Fig. 9), whose running sum closely recapitulates the saturated *a*-wave (Fig. 7A; 9C). The large, positive capacitive field potential (Fig. 9A, gray trace) is a biophysical consequence of the distribution of the current sources that charge the membrane capacitance during hyperpolarization (Robson and Frishman, 2014): a net decrease in outward current of the NKX and Kv2.1 channels in inner segment must occur for hyperpolarization of the cell body plasma membrane to occur: extracellularly, the consequence is a net increase in current flowing toward the inner segment. Were the corresponding capacitive field potential deleted (for example, if the capacitance were greatly reduced), there would be a much larger *a*-wave (Fig. 9B, C, gray traces), as the normal near-cancellation of the CNG field potential and capacitive field potential would be unbalanced in favor of the former.

Activation by hyperpolarization of the eponymous HCN1 channels necessarily occurs during the rod light response (Demontis et al., 2002), and the speed of this activation must be maximal in response to the strong stimulation that produces the saturated *a*-wave (Fig. S1). We previously developed a 2-state Boltzmann model of the rod HCN1 channels, and showed it provided a good description of the resting mouse rod I/V curve (Fortenbach et al., 2021), and that it to agreed reasonably well with measurements of HCN1 currents of rabbit rods (Demontis et al., 1999; Demontis et al., 2002; Demontis et al., 2009). While this HCN1 model may need further modification to provide a (perfect) description of the saturated rod photovoltage relaxation, when combined with the full rod model, the 2-state Boltzmann HCN1 model recovers three key kinetic features: the peak hyperpolarization, the plateau relaxation (~ −50 mV), and the approximate time constant of relaxation to the plateau (Fig. S2). (It bears emphasis, however, that in the living rod, voltage-dependent deactivation of Kv2.1 necessarily occurs during hyperpolarization and affects the time course of relaxation, both through its intrinsic kinetics and through increase in transmembrane resistance.) The result of deleting the trans-photoreceptor layer field potential of the HCN1 channels “in silico” is that the saturating *a*-wave is predicted to continue a slow negative-going trajectory, rather than relax toward the baseline (Fig. 9B, C).

The picture that emerges from our analysis is that the extracellular currents of both capacitive current and HCN1 current play ineluctable roles in determining the relaxation phase of the saturated *a*-wave. While it is clearly possible to produce a cable circuit model of the rod using resistive and capacitive components that neatly reproduces the *a*-wave relaxation Robson and Frishman, 2014), when the model rod incorporates all known ionic mechanisms (Fortenbach et al., 2021) with their measured axial distributions (Fig. 4), reproduction of the “nose” requires both components (Fig. 9C).

## SUPPLEMENTARY INFORMATION

**Figure S1.**
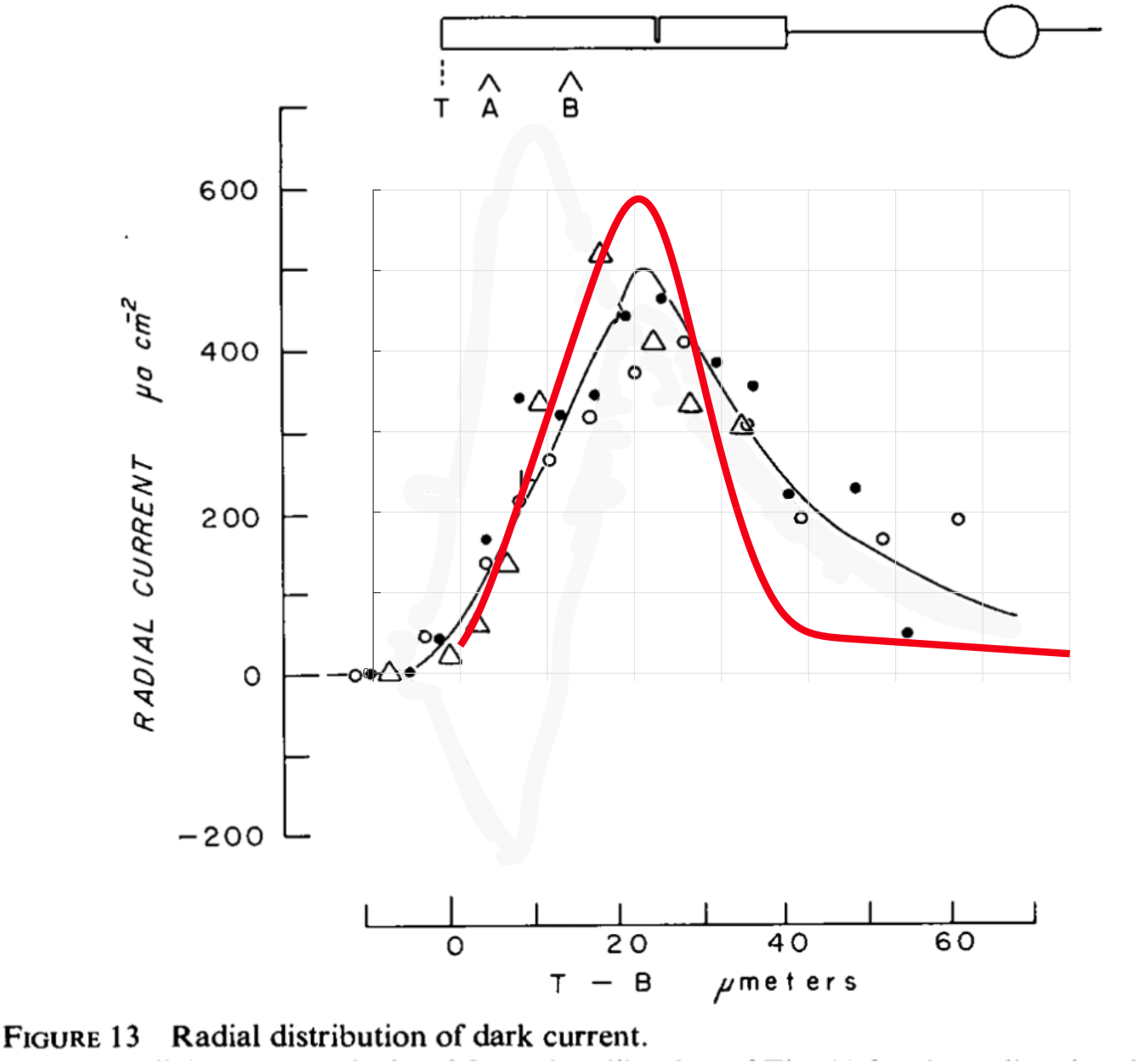
Ensemble model prediction of the rat rod layer current density measurements. The figure shows a screen shot of Fig. 13 of the classic results of Hagins, Penn & Yoshikami (1970) who discovered the rod dark current. The red curve is the current density profile generated by the rod ensemble model (cf. text Fig. 5D), and is presented on the same ordinate scale (gray grid) as the plot from Hagins et al,. The model has a rod density of 3.5×10^7^ rods cm^−2^ with individual rods having a dark current of 20 pA (in the outer segment this corresponds to a CNG-activated inward current of 19.3 pA and electrogenic Nckx inward current of 0.7 pA). The model curve was blurred with a Gaussian to reflect variation in the retinal of the rod inner segment (RIS)/rod outer segment (ROS) boundaries. The peak dark current density necessarily occurs at the ROS/RIS boundary, as in the rest state all current in the outer segment is inward, balanced by outward current primarily by NKX electrogenic current and Kv2.1 current in the inner segment (Fortenbach et al, 2021).

**Figure S2.**
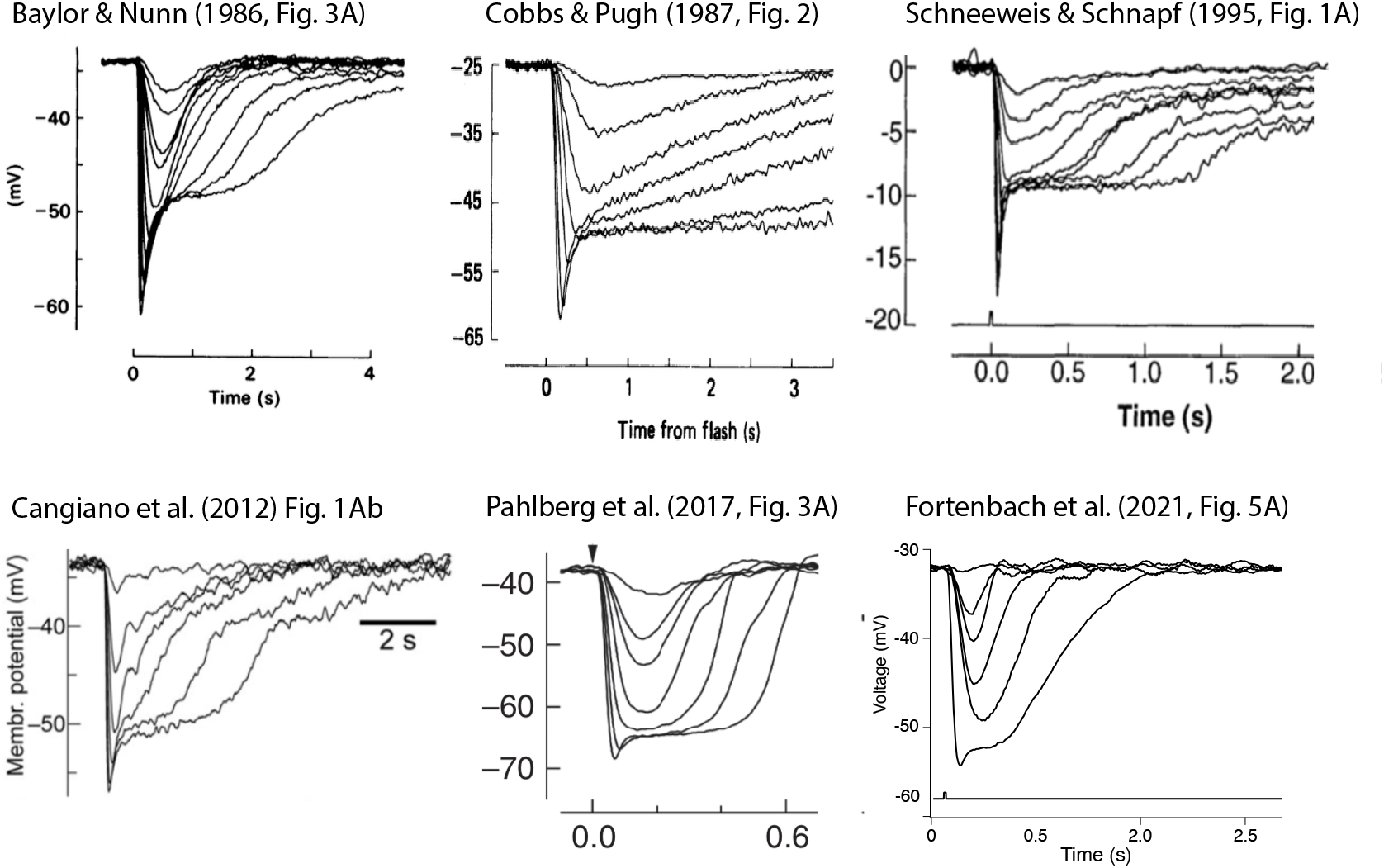
Published rod photovoltage response families. The source is indicated at the top of each figure, and provided in the notes to Table S1. The response families in the upper row were obtained from salamander rods (Baylor & Nunn, 1986) and Cobbs & Pugh (1987) and macaque (Schneeweis & Schnapf, 1995). All results in the lower row were obtained from mouse rods, but the temperature varied (cf, Table S1). The focus in the present investigation is on the saturated photovoltage, i.e., that which is obtained when the flash strength is so strong that the initial hyperpolarization velocity and peak are maximal. Responses in each family in the upper row appear to achieve saturation, but those in the lower row (particularly those of Pahlberg et al. and Fortenbach *et al*.) did not employ sufficiently stimuli sufficiently intense to produce saturated photovoltages.

**Figure S3.**
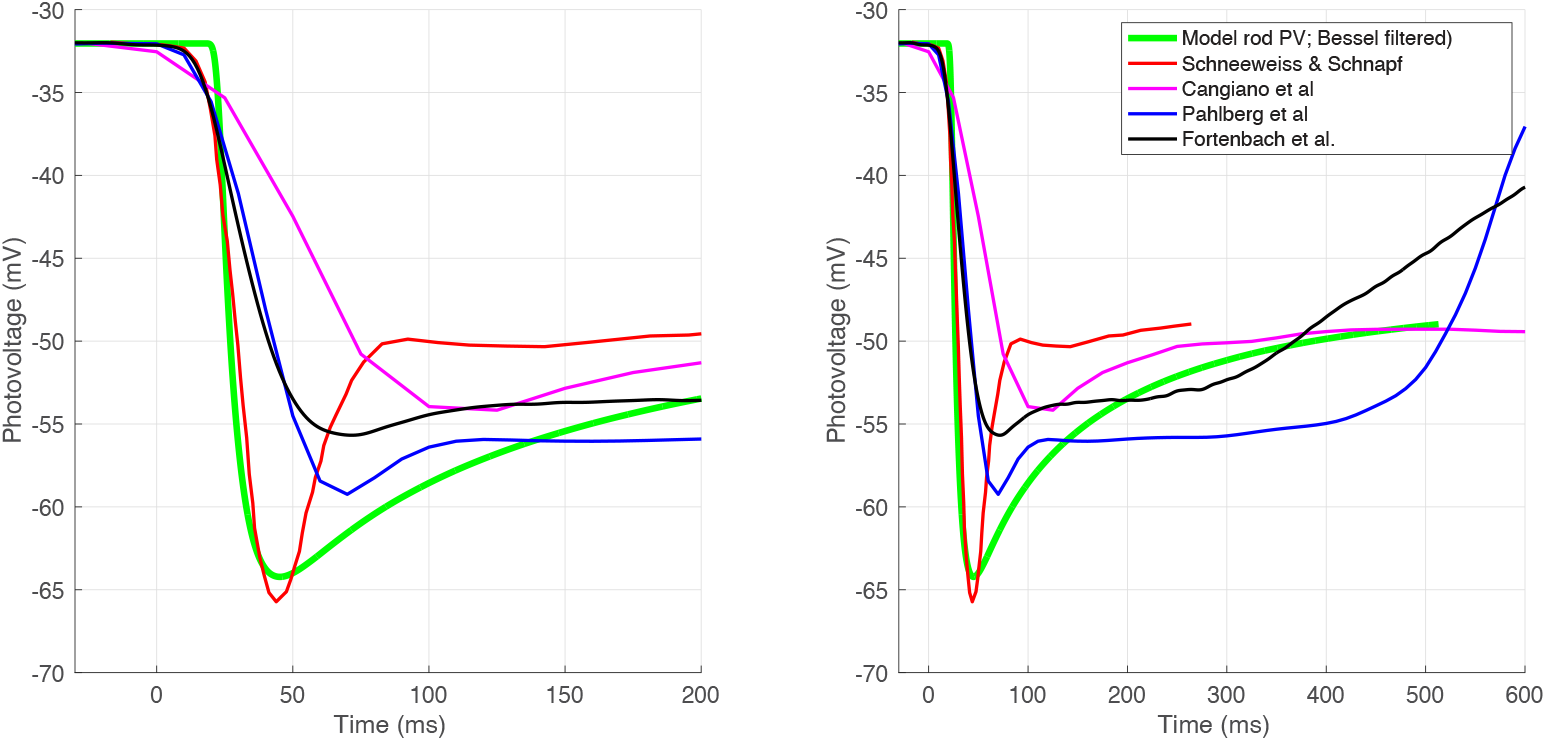
Rod photovoltage responses to the maximally intense stimuli used in the studies cited in Table S1. Both panels plot the same data, but the left-hand panel has a 3-fold expanded time scale to better illustrate key kinetic features. The data traces from other laboratories (red, magenta, blue) were extracted with the Matlab “Ungraph” kindly provided by T. D. Lamb, while that of Fortenbach *et al*. is the original digitized trace (cf. Fig. S2). The original trace from the study of Schneeweis & Schnapf (1995) had a zero level baseline (cf. Fig. S2): here it was offset to the resting potential of the mouse rod reported by Fortenbach *et al*. (2021) and scaled 1.9-fold, which brought the data into better correspondence with the other results. The trace from Cangiano et al. ((2012) The original trace from Pahlberg *et al*. (2017) was offset to the common resting potential, but otherwise not altered. The relaxation times in Table S1 were extracted from the traces in this figure.

### Prediction by Ensemble Model of the Current Density Measurements Hagins et al. (1970)

### Features of the photovoltage nose of rods, and rationalizing the use of the 2-state Boltzmann model of HCN1 gating

In response to strong stimulation the rod membrane photovoltage (PV) response relaxes from a maximally hyperpolarized state to a relatively depolarized plateau (Fig. S2). The saturated photovoltage response can be characterized by 4 parameters: the resting potential (*V*_m,rest_), the saturated peak hyperpolarization (Δ*V*_m,peak_), the plateau level (*V*_m,plat_), and the half-time of relaxation” (*t*_relax,0.5_) from the peak to the plateau. Figure S1 and Table S1 summarize the results from 5 studies, with the parameters extracted from the published data.

**Table S1:** Parameters of the Rod Photovoltage Response to Strong Flashes.

| Species | Temp | $V_{m,rest}$ | $V_{m,peak}$ | $V_{m,plat}$ | $\tau_{relax}$ | Reference |
| --- | --- | --- | --- | --- | --- | --- |
|  | C | mV | mV | mV | ms |  |
| Salamander | 20.4 | – 29 | – 62 | – 49 | 160 | 1, Fig. 3 |
| Salamander | 21 | – 25 | – 63 | – 50 | 150 |  |
| Macaque | 37 | – 37 | – 55* | – 47 | **42** | 2, Fig. 1; Fig S3 |
| Mouse | **24** | – 33 | – **57** | – 51 | (160) | 3, Fig. 1, Fig. S3 |
| Mouse | 36 | – 38 | (– 68) | – 64 | – | 4, Fig. 3, Fig. S3 |
| Mouse | 36 | – 32 | ( – 60) | – 53 | – | 5, Fig. 5, Fig. S3 |
| Model | – | – 32 | – 64 | – 49 | **130** | Fig. S3 |

*Table refs.* (1) Baylor & Nunn (1986); (2) Cobbs & Pugh (1987); (3) Schneeweis & Schnapf (1995); (4) Cangiano *et al.* (2012); (5) Pahlberg *et al.* (2017); (6) Fortenbach *et al.* (2021).
*Table note.* The lowermost row of the table applies to the 2-state Boltzmann model of HCN1 gating of Fortenbach *et al.* (2021), which is used in this investigation without modification.

The relaxation of *V*_m_ from the peak hyperpolarization to the plateau is generally thought to be caused by the activation of HCN1 channels: thus, rapid and sustained closure of all CNG channels causes *V*_m_ to hyperpolarize at towards E_K_, the reversible potential of the K_v_2.1 channels that were open in the relatively depolarized dark state, and the hyperpolarization activates the eponymous HCN1 channels. For sufficiently intense flashes, the CNG channels close in a few milliseconds, and the maximum rate of hyperpolarization is limited by the membrane time constant, dictated primarily by the resting K^+^ conductance and of course capacitance (Cobbs & Pugh, 1987). The most critical parameter for the present investigation is the relaxation time from peak to plateau, as the question being addressed is whether the nose of the *a*-wave is determined by extracellular flow of capacitance current or by extracellular flow of HCN1 current. To extract the HCN1 activation time, we approximated the relaxations with an exponential decay (*τ*_relax_, column 5 of Table S1) of the photovoltage responses from the above studies (the only ones of which we are aware that have PV families). Observance of the HCN1 relaxation from maximal hyperpolarization requires exposure to very strong flashes, which were not used in the studies by Pahlberg et al. and Fortenbach et al‥ Notably, the HCN1 activation appears sensitive to temperature, as seen in the comparison of salamander at 20-21C, mouse at 24C, and macaque at 37C.

Two-state Boltzmann models of Kv2.1and HCN1 gating were presented Fortenbach *et al*. (2021) and used primarily to explain the dark adapted mouse rod I/V curve (which exhibits a very strong inward current at hyperpolarized potentials). However, they were also used in that paper and herein for kinetic modeling the mouse rod photovoltage response to very strong light stimuli. To evaluate its utility for this latter purpose we present data extracted from the published mouse rod photovoltage responses to the maximally intense stimuli used by different investigators and the results from a macaque rod (Schneeweis & Schnapf, 1995) (Fig. S3). The saturated photovoltage response of the model of Fortenbach *et al*. (2021) (Fig. S3, green trace) has a plausible peak (− 64 mV) and relaxes to a reasonable plateau level (− 49 mV). However, in comparison with the scaled version of the macaque saturated photovoltage response, the relaxation time is slow: 130 ms vs. 42 ms.

While this analysis suggests the 2-state kinetic models of Kv2.1 and HCN1 may need revision in the future when truly saturated mouse rod photovoltage data are available, it bears emphasis that they were developed primarily to explain the resting I/V curve of the mouse rod and the earliest (hyperpolarizing phase) of the mouse rod photovoltage. It also should be emphasized that the underlying model of phototransduction used here involves no inactivation of the transduction cascade in the 50 ms time frame (Figs. 6-9) applicable the *a*-waves of rods stimulated with intense flashes.

